# Tunable Rigid Spikes on Virus-Like Porous Silica Enable Mechanistically Controlled Nanovaccine Platforms

**DOI:** 10.64898/2026.04.26.720861

**Authors:** Cui Pang, Jiajia Wang, Ahmed Montaser, Shuhao Ma, Henri Leinonen, Guoqing Hu, Vesa-Pekka Lehto, Li Fan, Wujun Xu

## Abstract

Virus-like particles represent an emerging and promising vaccine platform. However, these particles are inherently mechanically soft and have limited control over particle surface architecture, thereby constraining their immunological control. Herein, we report the rational design of bioinspired virus-like porous silica (VLPSi) nanoparticles (NPs) with tunable and mechanically rigid spike architectures that function dually as antigen delivery carriers and immune adjuvants. Using ovalbumin (OVA) as a model antigen, we systematically elucidate the spiky structure–function relationship in antigen delivery and immune response. VLPSi NPs exhibit good biocompatibility, sustained antigen release, and markedly enhanced cellular uptake and endosomal escape compared with soft spike and spherical counterparts. Mechanistic investigations combining molecular dynamics simulations and proteomic analyses reveal that rigid spike architectures reduce the energetic barrier for cellular internalization and concurrently activate dual pathways involving endosomal Toll like receptors and calcium signaling. Consequently, VLPSi with long spikes elicit significantly enhanced humoral and cellular immune responses, outperforming the particles with shorter spikes, spherical shape as well as clinically used alum adjuvant. To demonstrate translational potential, bioinspired antibacterial vaccines were produced by loading Staphylococcus aureus surface protein rEsxB. The VLPSi-based vaccine elicited robust protective immunity to achieve complete (100%) survival following lethal challenge without detectable adverse effects, whereas traditional Alum-adjuvanted formulation conferred only minimal protection, with a survival rate of 10%. Collectively, this work establishes VLPSi with tunable spikes as a mechanistically controlled platform for next generation vaccines.

**Graphic Abstract:** 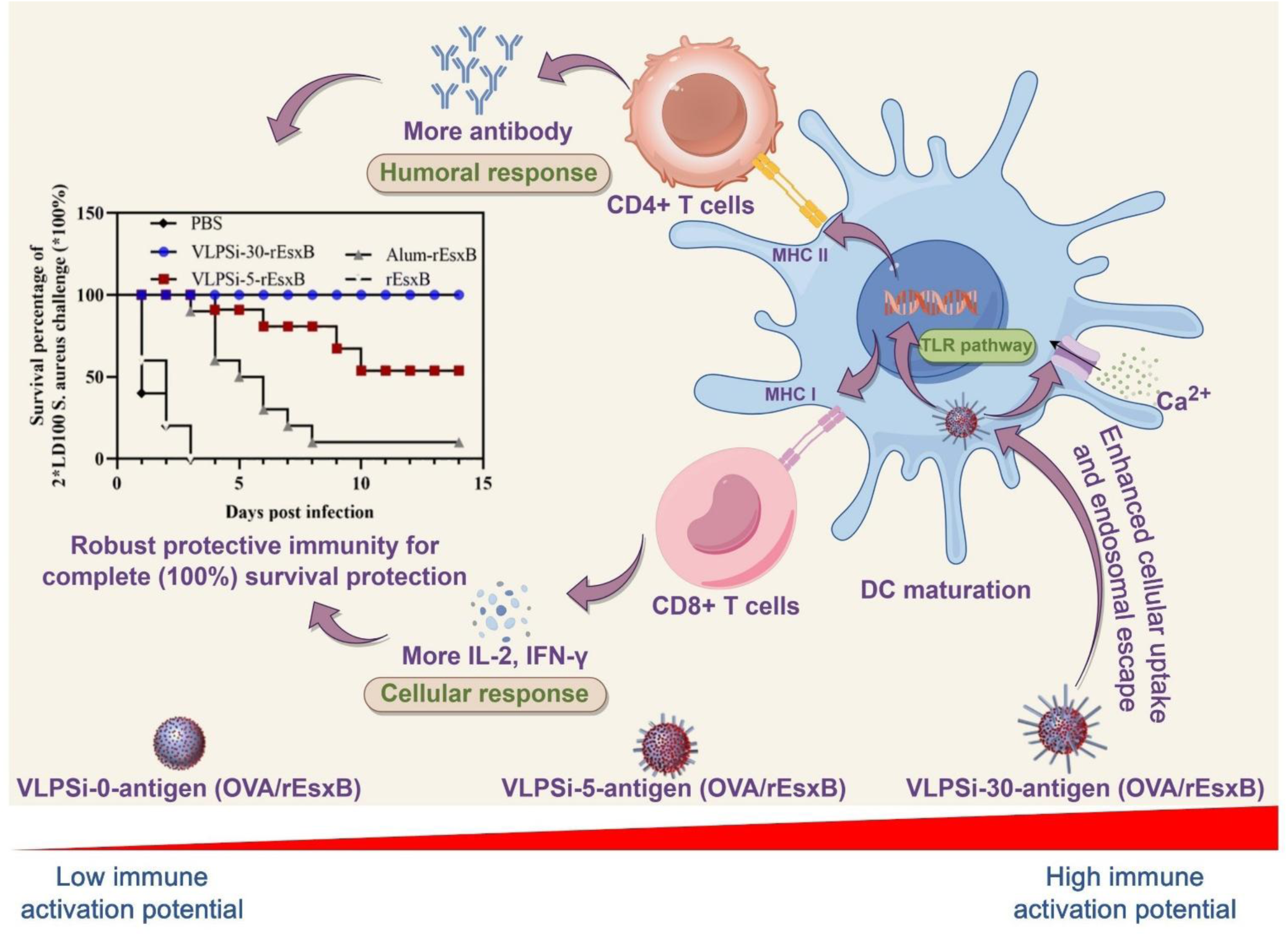

## Introduction

Vaccines are a highly effective and safe strategy for preventing infectious diseases by triggering the host’s immunity against specific pathogens^1^. Compared to anti-bactericidal drugs such as antibiotics, anti-bactericidal vaccines reduce the emergence of drug-resistant strains and superbugs. Vaccines typically consist of both antigen and adjuvant, with the latter to promote the immune response and antibody production. Adjuvants can be classified into immunostimulatory and antigen-delivery categories based on their mechanisms of immune enhancement. Immunostimulatory adjuvants, such as Toll-like receptor (TLR) agonists, act as pathogen-associated molecular patterns (PAMPs), damage-associated molecular patterns (DAMPs), or their mimics to react with pattern recognition receptors (PRRs) on antigen-presenting cells (APCs). This interaction initiates an innate immune response that directly facilitates the activation and maturation of APCs for antigen presentation^2^. However, delivery system is generally required for most immunostimulatory adjuvants. On the other hand, antigen-delivery adjuvants which serve as both self-adjuvants and carriers to load antigens, thereby co-locally augmenting the uptake and presentation of antigens by APCs^3^.

Dendritic cells (DCs) are important types of APCs which play a pivotal role in evoking innate and adaptive immune responses^4^. Once activated, they migrate to lymphoid tissues, present antigens to naïve T cells, and evoke downstream cellular and humoral immunity effectively. Therefore, targeted activation of DCs is critical for determining vaccine efficacy, especially for inducing robust and long-lasting protection. Recent research has explored nanoparticles (NPs)-based strategies to develop antigen-delivery adjuvants for activating DCs. Different types of NPs, such as polymers^5^, proteins^6^, lipids^7^, gold^8^, mesoporous silica^9^ have been tailored with surface ligands or charge modifications to improve DC activation. These advances highlight the potential of nanotechnology to design novel immune adjuvants with effective DC modulation. Nevertheless, specific chemical compositions or surface chemical ligands are the main reasons to endow immune adjuvant property on these NPs.

The morphology of NPs is an important factor affecting immune activation. Virus-like particles have been widely used as both adjuvants and antigens delivery system for vaccines^10, 11^. However, up to now, virus-like NPs were generally prepared by assembling viral proteins, which are physically ‘soft’. Previous study revealed that particle rigidity affects the effectiveness in cell uptake and activating APCs. Soft NPs are easily deformed by the force originating from cell membrane wrapping, which reduces the cellular binding for decreased cell uptake^12^. Furthermore, it was found that rigid particles demonstrated >100 times greater efficiency in priming DCs in draining lymph nodes than “soft” counterparts^13^. TiO_2_ particles with rigid nanospikes have shown the potent capacity to activate and amplify immune responses both *in vitro* and *in vivo* ^14^. These spiked particles have been observed to enhance antigen-specific humoral and cellular immune responses, thereby stimulating protective immunity against tumor growth and influenza virus infection. This study offers valuable insights into the role of rigid spikes in regulating the activation of innate immunity ^14^. However, TiO_2_ is hardly degradable in biological fluids. Alternatively, biodegradable silica-based virus-like particles have been developed for as effective adjuvants for vaccines^11^. Li et al developed raspberry-like silica-adjuvanted nanovaccines by coating virus-like viral particles with spherical silica^11^. The nanovaccines enhance the secretion of both Th1 and Th2 type cytokines in murine bone marrow-derived dendritic cells (BMDCs) and improve DC activation in draining lymph nodes. However, to the best of our knowledge, the application of virus-like porous silica (VLPSi) NPs with tunable rigid spikes as immune adjuvants has not yet been investigated. Consequently, significant knowledge gaps remain regarding their immunological efficacy and the mechanisms by which they activate DCs.

In the present study, we aim to demonstrate the application of VLPSi as novel DC-activating adjuvants by tuning the length of surface spike. First, VLPSi NPs with the spike length of 0 (VLPSi-0), 5 (VLPSi-5), 30 nm (VLPSi-30) were synthesized. OVA was chosen as the model antigens in the initial proof-of-concept study (Scheme 1). The mechanism in activating immune response of VLPSi was explored by using computation simulation and proteomics. To further verify the potential of VLPSi adjuvants, we developed an antibacterial vaccine by conjugating VLPSi with the antigen of rEsxB, a recombinant protein from Staphylococcus aureus. The performance the antibacterial vaccine, including antibody expression, cytokines, mice survival in lethal challenge, and safety, were thoroughly assessed *in vivo* (Scheme 1).

**Schema 1.**
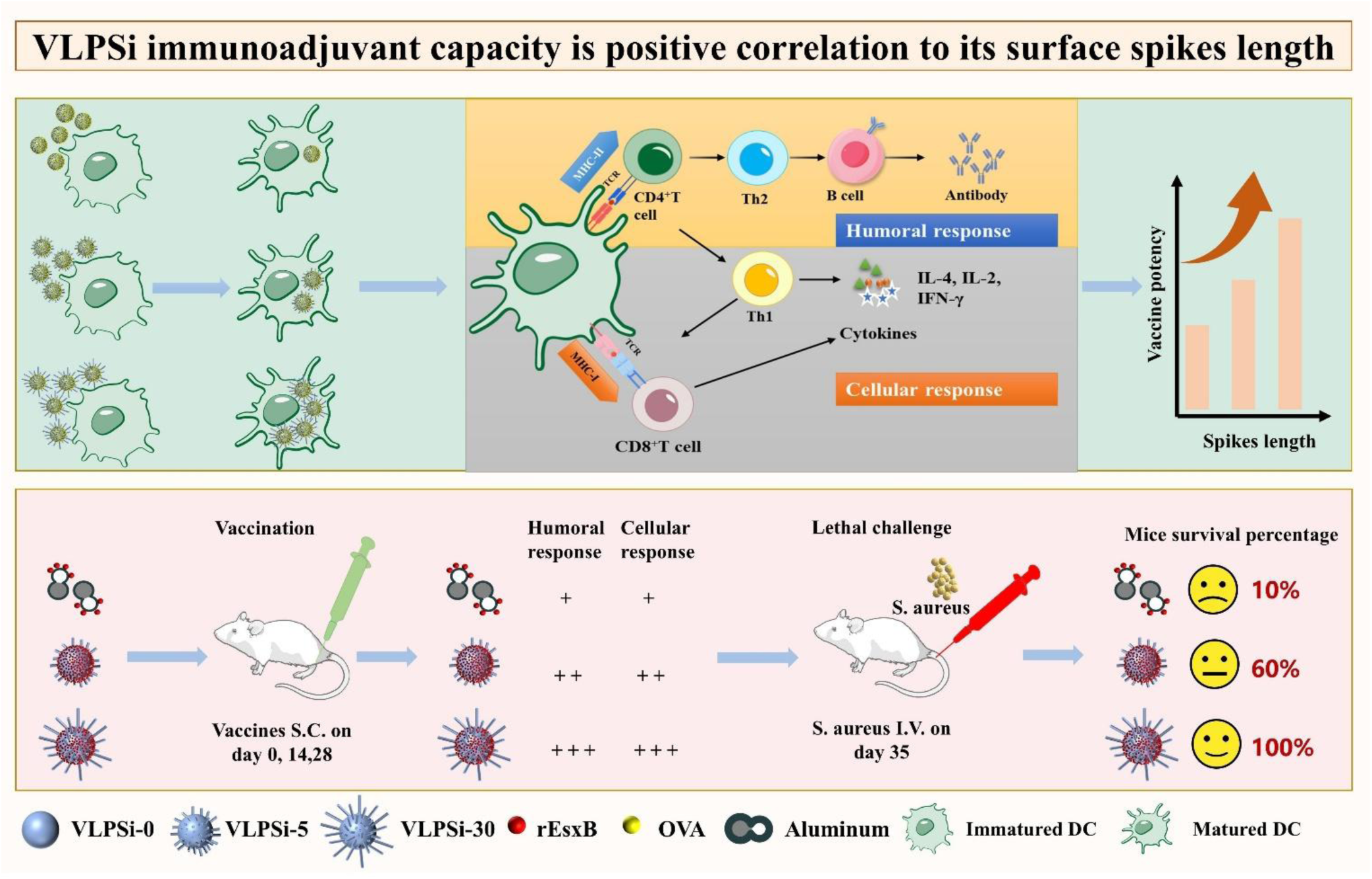
Schematic representation of the preparation of VLPSi and the exploration of its immune activation ability. The effect of different VLPSi on the activation capacity of BMDCs was first examined using the OVA antigen. Subsequently, S. aureus rEsxB was loaded to verify the stronger immune activation and protection provided by VLPSi-30 in antimicrobial vaccines. Longer surface VLPSi enhanced the phagocytosis of DCs, promoted their maturation, and increased the secretion of interleukin-2 (IL-2), interleukin-4 (IL-4) and interferon (IFN-γ), thereby enhancing immune activation in mice.

## 2. Experimental Methods

### 2.1 Materials

The following chemical reagents were purchased and used directly: hexadecyltrimethylammonium bromide (CTAB, Sigma), Tetraethyl orthosilicate (99% GC, TEOS, Merck), Cyclohexane (Merck), 0.1 M Sodium hydroxide, sodium acetate trihydrate, hydrochloric acid, 2-[Methoxy(polyethyleneoxy)propyl] trimethoxy silane (90%, 6-9 PEG-units, abcr), 3-Aminopropyltriethoxysilane (APTES, VWR), Electron Microscopy Sciences (HC400-Cu, USA), Albumin from chicken egg white (OVA, Sigma), The Pierce BCA Protein Assay Kit (Fisher Scientific), Phosphate buffer saline (PBS, VWR), fetal bovine serum (FBS, Thermo Fisher Scientific), Dulbecco’s Modified Eagle Medium (DMEM, Thermo Fisher Scientific), Hank’s balanced salt buffer(HBSS, Thermo Fisher Scientific), Cell Titer-Glo® 2.0 Cell Viability Assay (Promega), Fluorescein 5(6)-isothiocyanate(FITC, Alfa Chemical), Invitrogen CellMask™ plasma membrane stains (Thermo Fisher Scientific), 4’,6-Diamidino-2-Phenylindole(DAPI, Sigma), Cell Extraction Buffer PTR (ab193970) and Cell Extraction Enhancer Solution (ab193971), low-protein binding tubes (Thermo Fisher Scientific), Dithiothreitol (DTT) (Merk), Iodoacetamide (Merk), Sera-Mag magnetic carboxylate beads (Cytiva, 24152105050250), Sera-Mag magnetic carboxylate beads (Cytiva, 44152105050250), Ethanol (Sigma), Pierce Trypsin Protease (Thermo Fischer), thermomixer (Thermo Fischer), SpeedVac vacuum concentrator (Thermo Fisher), Agilent AdvanceBio Peptide Map column (2.1 mm × 250 mm, 2.7 μm, Agilent Technologies), Ultrapure Milli-Q water (18.2 MΩ cm, Millipore). BL21 (DE3)-competent Escherichia coli was procured from TIANGEN BIOTECH (BEIJING) Co., Ltd. (Beijing, China). Ni-Sepharose was obtained from Cytiva (Shanghai, China), and a High-Capacity Endotoxin Removal column was acquired from Xiamen Bioendo Technology Co., Ltd. (Xiamen, China). Various reagents, including isopropyl β-D-thiogalactopyranoside (IPTG), imidazole, acetone, 2-(N-morpholino)ethanesulfonic acid hydrate (MES), 1-ethyl-3-(3-dimethylaminopropyl)carbodiimide (EDC), N-hydroxysuccinimide (NHS), bovine serum albumin (BSA), Tween 20, 2,2’-azino-bis(3-ethylbenzothiazoline-6-sulfonic acid) (ABTS), horseradish peroxidase (HRP)-labeled goat anti-mouse IgG, and 0.22 μm and 0.45 μm sterile filters were sourced from Merck (Shanghai, China). Mouse IL-4 ELISA Kit (NBS EMC002.96), Mouse IFN-γ ELISA Kit (NBS EMC101g.96), Mouse IFN-γ ELISpotSASIC (HRP) (Mabtech 3321-2H), Mouse IL-4 ELISpotSASIC (HRP) (Mabtech 3311-2H), Tetramethylbenzidine (TMB) substrate for ELISpot (Mabtech 3651-10), OVA IgG ELISA Kit (Chondrex 3011), OVA IgM ELISA Kit (Chondrex 3017), and Low Endotoxin Ovalbumin from Chick Egg White (Chondrex 3022).

### 2.2 Methods

#### 2.2.1 Preparation of VLPSi NPs

VLPSi NPs were prepared according to the modified protocol published before^15^. CTAB was dissolved in Ultrapure Milli-Q water in a round-bottom flask to prepare the surfactant solution with 1% CTAB, and then 0.1 M NaOH solution was slowly added. Next, TEOS with 20 % mass concentration in cyclohexane was slowly added into solution. The VLPSi with different spiky lengths were obtained after the reaction at 60 °C for 24-72 h. Finally, the VLPSi was washed with ethanol twice and refluxed with ethanol-hydrochloric acid mixture for 48 h to remove the surfactant template.

Smooth NPs (VLPSi-0, referring to 0 nm spikes on the surface of NPs) were prepared as reference samples. Specifically, 6.24 g CTAB was dissolved in 53.4 mL Ultrapure Milli-Q water and then 0.3 g sodium acetate trihydrate was added. 4.35 mL TEOS was slowly added into the mixed solution and stirred at 60 °C for 24 h. The sample was washed with ethanol twice, followed by refluxing in a mixture of ethanol and hydrochloric acid for 48 h. The final samples were stored in ethanol for further use.

#### 2.2.2 Surface modifications and characterization of NPs

(1) PEGylation of VLPSi. The obtained VLPSi (10 mg) were dispersed in 2 mL of ethanol and mixed with 200 uL PEG-silane (0.5 kDa) ^16^. After sonicated for 2 min, the mixture was stirred and bubbled under air atmosphere to evaporate ethanol at 90 °C for 30 mins. The reaction continued at the same temperature for 30 mins. Finally, the product was redispersed in 2.0 mL ethanol and then washed twice with ethanol. The obtained VLPSi was noted as PEG-VLPSi.

To study the effect of spike stiffness on cell uptake and endosomal escape, NPs with soft spikes were constructed by grafting two different lengths of PEGs. Thus, 0.5 and 2.0 kDa PEG-silanes were mixed in toluene at first. VLPSi-0 and VLPSi-30 were added into the mixture and reacted at 90 °C for overnight. The NPs were centrifugated and washed with ethanol twice.

(2) Amine modified VLPSi. 10 mg PEG-VLPSi was dispersed in 2 mL of ethanol. Then, 3 µL APTES were added, and the mixture was stirred at 65°C for 1 h. The samples were collected by centrifuging and washed twice with ethanol. The obtained samples were noted as PEG-VLPSi-NH_2_.

(3) Characterization of VLPSi. The VLPSi were dispersed in ethanol by ultrasound for 5 mins, and then one drop of the solution was left on the copper grid. After dried at room temperature, the samples were imaged with transmission electron microscopy (TEM) (JEOL JEM-2100F). The surface charge of the VLPSi was measured with Zetasizer (Malvern Panalytical Ltd, United Kingdom).

#### 2.2.3 OVA loading and release

8 mg OVA was dispersed in 1 mL PBS and mixed with 10 mg PEG-VLPSi-NH_2_ with different spike lengths (0, 5, 30 nm), and the solution was stirred at room temperature overnight. The collected samples were noted as PEG-VLPSi-NH_2_-OVA-0, PEG-VLPSi-NH_2_-OVA-5, and PEG-VLPSi-NH_2_-OVA-30, which are abbreviated as VLPSi-0-OVA, VLPSi-5-OVA, and VLPSi-30-OVA, respectively. Then, the concentration of free OVA in supernatant was measured by the Pierce BCA Protein Assay Kit. The loading efficiency of OVA was calculated according to the following equation: loading efficiency (%) = (total OVA mass - OVA mass in supernatant)/total OVA mass × 100%.

To evaluate the OVA release, the VLPSi-0-OVA was dispersed in 1 mL PBS with a rotating mixer at 37°C. Supernatant was taken with centrifuge at 1, 2, 4, 6, 8, and 24 h. The released OVA concentration was measured according to Pierce BCA Protein Assay Kit. All experiments were performed in triplicate.

#### 2.2.4 Safety of NPs

BALB/C mice were obtained from the Animal Experimentation Center of the Air Force Medical University. All procedures complied with the National Research Council’s guidelines and were approved by the Animal Care and Ethics Committee of the Fourth Military Medical University (Approval No. KY20213144-1). Six-week-old BALB/c mice were used to aseptically isolate bone marrow. After centrifugation and erythrocyte lysis, the cell pellet was resuspended in RPMI-1640 complete medium and adjusted to 1×10^6^ cells/mL. Recombinant Mouse GM-CSF was added to a final concentration of 10 ng/mL, and cells were cultured at 37 °C in a 5% CO_2_ incubator. On day 3, fresh medium containing GM-CSF was replenished. By day 7, suspended and semi-suspended cells were collected as bone marrow-derived dendritic cells (BMDCs).

Macrophages, Human Umbilical Vein Endothelial Cells (HUVECs) and mouse BMDCs were utilized to assess the cytotoxicity of VLPSi-0-OVA, VLPSi-5-OVA, and VLPSi-30-OVA. Cells were seeded in 96-well plates at a density of 5×10^3^ cells per well and allowed to adhere overnight. On the following day, NPs were introduced into the cell culture medium at final concentrations of 25 μg/mL, 50 μg/mL, 75 μg/mL, 100 μg/mL, and 200 μg/mL respectively. Each sample was prepared in six replicate wells. The cells were subsequently incubated for an additional 24 hours. Thereafter, Cell Counting Kit-8 (CCK-8) solution was added to each well to achieve a final concentration of 10%, and the plates were incubated for 2 hours at 37°C. Optical density (OD) was measured at 450 nm using a microplate reader (Biotek SYNERGY LX Instruments, Santa Clara, CA, USA). Cell viability was calculated using the standard cytotoxicity equation: Cell Viability = [(OD_test_ - OD_background_) / (OD_control_ - OD_background_)] × 100%.

#### 2.2.5 Coarse-grained molecular dynamics simulation

To investigate the interaction between nanoparticles and cell membranes or lysosomes, we use the solvent-free coarse-grained model developed by Cooke and Deserno^17^. This model has been widely validated for studying membrane–NP interactions^18^.

For NP - membrane interactions, the lipid bilayer is constructed with a box dimension of 210 σ × 210 σ × 500 σ, and the cell membrane is modeled as a periodic planar membrane. In our simulations, each lipid molecule is represented by three connected beads: one for the hydrophilic head and two for the hydrophobic tail. The repulsive interaction between beads is modeled using the Weeks–Chandler–Anderson potential:

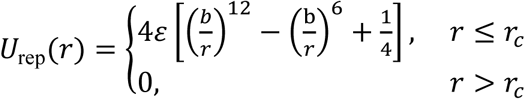

where 𝑏_ℎ𝑒𝑎𝑑,ℎ𝑒𝑎𝑑_ = 𝑏_ℎ𝑒𝑎𝑑,𝑡𝑎𝑖𝑙_ = 𝑏_ℎ𝑒𝑎𝑑,𝑁𝑃_ = 0.95σ, 𝑏_𝑡𝑎𝑖𝑙,𝑡𝑎𝑖𝑙_ = 𝑏_𝑡𝑎𝑖𝑙,𝑁𝑃_ = 1.0σ, 𝑟_𝑐_ = 2.5𝑏, and 𝑟 is the distance between beads. An attractive potential is applied between tail-tail, and tail-NP to mimic hydrophobic interactions:

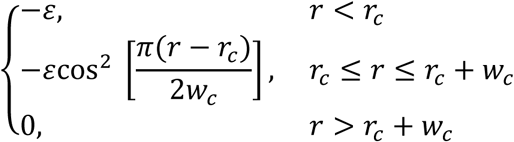

where 𝑟_𝑐_ = 2^1/6^𝑏, 𝑤_𝑐_ = 1.6σ. Bonds between adjacent beads within a lipid molecule are modeled using the finite extensible nonlinear elastic (FENE) potential:

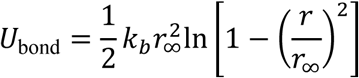

where 𝑘_𝑏_ = 30𝜀, 𝑟_∞_ = 1.5σ. To account for lipid stiffness, a harmonic bending potential is applied between the head bead and the second tail bead:

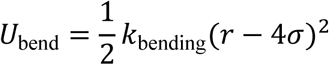

where 𝑘_bending_ = 10𝜀.

Since lysosomes are comparable in size to NPs, lysosomes are modeled as three-dimensional structures rather than planar membranes. The lysosomes are first modeled as 𝑁_t_ triangular networks composed of 𝑁 beads as the innermost beads of lysosomes, and then extend outward along the normal direction to develop a phospholipid bilayer model. Considering the incompressibility of the liquid inside the lysosome and the weak elasticity of the lipid membrane, the elastic energy of the surface area and volume are introduced^19^

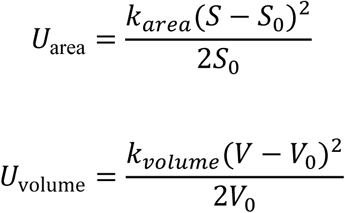

where 𝑆 and 𝑉 are the surface area and volume of the lysosomes. Subscript 0 represents the target value, and 𝑘_𝑎𝑟𝑒𝑎_ = 5000𝜀, 𝑘_𝑣𝑜𝑙𝑢𝑚𝑒_ = 500𝜀 are the corresponding constraint coefficients. For NP – lysosomes interactions, additional Morse potential id applied to mimic the phagocytosis of lysosomes on NPs^20^

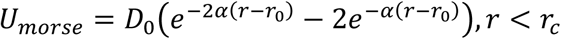

where 𝐷_0_ = 60𝜀, 𝛼 = 0.5σ^−1^, 𝑟_𝑐_ = 2.0σ.

All simulations are conducted using LAMMPS^21^. The time step is set to Δ𝑡 = 0.002𝜏, where the time unit 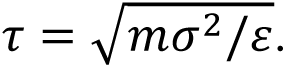 The system temperature is maintained at 𝑘_B_𝑇 = 1.1𝜀 using a Langevin thermostat with a damping constant of 2.0 𝜏. A modified Berendsen barostat^22^ is applied to maintain zero membrane tension with a time constant of 2.0 𝜏 for the membrane case, while the case of lysosomes adopted the NVT ensemble. Periodic boundary conditions are applied in all three directions. Comparing to a typical membrane thickness 𝑡 = 10 nm, the bead diameter, 𝜎, is set at 2 nm. The mass of the beads is all set to 1 in our model, and the time unit 𝜏 to 40 ns. it takes ∼5.0 × 10^6^Δ𝑡 for a typical simulation performed in the current study. The spiky NPs consist of a sphere from which cylindrical spikes grow outward. The radius of the sphere is 𝑟 = 30 nm, while each cylinder has a radius of 𝑎 = 5 nm and a length of 𝐿, and the surface density of the spikes is 𝜌 = 0.0018 nm^−2^. The number density of the CG beads that make up each NP is set to 4.

To thermodynamically illustrate the important role of spike in the interaction between NPs and cell membranes, we calculate the changes of free energy of NPs as they cross the cell membranes. A spring force with a constant of 2000𝜀 is applied to the center of mass of the NP at a slow rate of 0.005𝜎/𝜏. Based on the principle of Jarzynski equality^23^

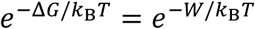

where Δ𝐺 denotes the free energy change, and 𝑊 represents the work exerted on the moving the NPs cross the cell membranes.

The lysosomal rupture simulations start with a lysosome capable of engulfing a nanoparticle, with 𝑆 = 1.75 × 10^6^ nm^2^ and 𝑉 = 3.86 × 10^7^ nm^3^. This is obtained by a lysosome close to a given NP simulated by an equilibrium state of about 50,000,000 steps. The relative positions of lysosomes and nanoparticles are constrained to simulate the rupture of lysosomes.

### Theoretical model

Inspired by the theoretical description of limiting geometry conditions of cell flow^24^, a simplified axisymmetric model (Schema 2) is considered to clarify how geometric constraints affect phagocytosis of nanoparticles by lysosomes. We assume that after a given volume of lysosomes completely engulfs a nanoparticle, in the limiting case, the remaining part remains spherical, which minimizes the required surface area of the lysosomes. Ignoring the elastic deformation of the volume and surface area of lysosomes, the surface area and volume of lysosomes can be expressed as

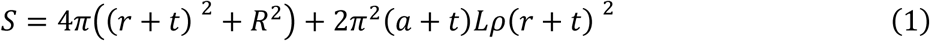

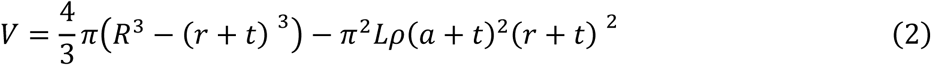

where the membrane thickness 𝑡 is considered non-negligible. From Eq. (2), 𝑅 can be expressed as

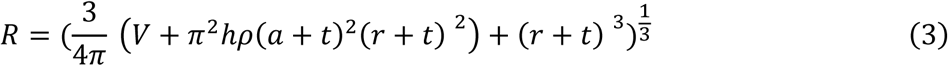

With the substitution of Eq. (3) into (1), one can obtain the relationship between the surface area 𝑆 and the volume 𝑉, which is given by

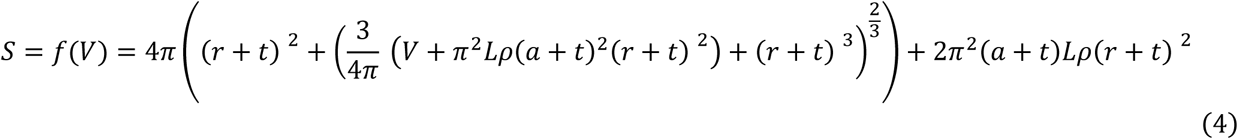

**Schema 2.**
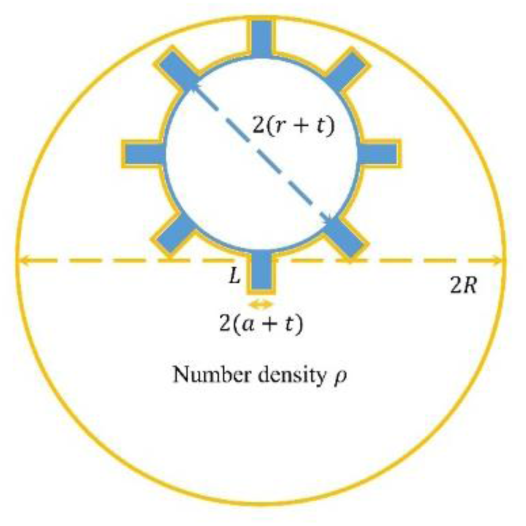
Schematic diagram of the limiting geometry considered in the theoretical model for lysosomal engulfing nanoparticles. The yellow lines represent the lysosome membrane, and the blue part denotes the NP. The model is considered axisymmetric.

#### 2.2.6 Cellular internalization

The VLPSi with different spike lengths were labeled with fluorescence dye FITC before internalization experiments^25^. 10 mg of PEG-VLPSi-NH_2_ was mixed with 1 mg of FITC in 2 mL of phosphate buffered saline (PBS). The mixture was stirred for 12 h at room temperature in dark conditions. The FITC-labelled samples were noted as FITC-VLPSi-0-OVA, FITC-VLPSi-5-OVA, and FITC-VLPSi-30-OVA.

RAW 264.7 cells were divided into the 8-well plate at a density of 2х10^4^ cells per well. The cells were cultured in DMEM with 10 % fetal bovine serum (FBS) for 24 h at 37 °C and 5% CO_2_ atmosphere. The FITC-VLPSi-OVA were incubated with the cells at a concentration of 50 µg/mL. After 4 h, the cells were washed with 200 µL HBSS three times to remove non-internalized particles. The cell membranes were labeled with 5 µg/mL CellMask™ red plasma membrane stain for 10 min and then fixed with 4% paraformaldehyde solution. After that, the cell nuclei were stained with 10 µg/ml DAPI solution for 10min and then washed by 200 µL HBSS twice. The cell internalization was imaged with a confocal laser scanning microscopy (Zeiss LSM 700). The channels of blue (DAPI), green (FITC), red (CellMask™) were observed using 405, 488, and 550 nm laser excitation, respectively. To study endosomal escape of the NPs, cells were incubated with the similar protocol as described above, except cells were stained with lysosome probe LysoTracker™ Red (Invitrogen).

#### 2.2.7 Flow cytometry

The NPs were dispersed in 1.0 mL of PBS solution and rotated for a few seconds. Free OVA was used as reference sample. Each sample was diluted to predetermined concentrations with cell medium. BMDCs were seeded in 12-well plates at the density of 1 × 10^6^ cells per well. Subsequently, OVA loaded NPs were added to the wells and incubate for 24 h. Next, the cells were detached from the plates and washed three times with PBS. The cells were then analyzed with flow cytometry (NovoCyteTM and Agilent) to assess the activation markers (CD 80, CD 86, and CD 40) of DCs. CD40 was conjugated with the fluorescent dye phycoerythrin (PE), CD80 with fluorescein isothiocyanate (FITC), and CD86 with PE-Cy7. The expression levels and mean fluorescence intensities (MFI) of markers, including CD40, CD80, and CD86, were analyzed using FlowJo V.10.8.1. Graphs were generated using GraphPad Prism.

#### 2.2.8 Effect of VLPSi-OVA on cytokine secretion by BMDCs

BMDCs were seeded in 48-well plates at a density of 5×10^4^ cells per well. According to the experimental groups, 500 µL of RPMI 1640 complete medium (supplemented with 10% FBS), free OVA, and OVA loaded NPs were added to the wells and incubated for 18 hours. Following incubation, the cell culture supernatant was aspirated and centrifuged at 300 g for 5 minutes. The supernatant was then subjected to enzyme-linked immunosorbent assay (ELISA) for analysis. A series of interleukin-1 beta (IL-1β) standards were prepared at concentrations of 500, 250, 125, 62.5, 31.25, 15.6, 7.8, and 0 pg/mL. To the blank wells, add the universal diluent, and to the remaining corresponding wells, introduce varying concentrations of standards (100 µl per well). Incubate the plate at 37°C for 90 minutes, ensuring it is protected from light. Subsequently, wash the plate five times. The blank wells should then be treated with biotinylated antibody diluent, while the remaining wells receive the biotinylated antibody working solution (100 µl per well). Incubate again at 37°C for 60 minutes, maintaining light protection, and wash the plate five times. Next, spike the blank wells with enzyme conjugate diluent and the other wells with enzyme conjugate working solution (100 µl per well). Conduct incubation at 37°C for 30 minutes, ensuring protection from light, followed by five washes of the plate. Introduce the color development substrate (TMB) at 100 µl per well and incubate at 37°C for 15 minutes. Add 100 µl per well of the reaction termination solution, mix thoroughly, and immediately measure the optical density at 450 nm. The concentrations of IL-6 and IFN-γ were determined following the previous published protocol^26^.

#### 2.2.9 Proteomics

RAW 264.7 cells (6×10⁴ per well) were seeded in six-well plates, incubated for 24 hours, and treated with VLPSi-0-OVA, VLPSi-5-OVA, and VLPSi-30-OVA nanoparticles (50 µg/mL) for 12 hours. Cells were washed with cold PBS, lysed with 200 µL of ice-cold Cell Extraction Buffer for 15 minutes, and centrifuged at 18,000 g for 20 minutes. The supernatant, containing solubilized proteins, was collected in low-protein-binding tubes. Protein quantification was performed using the Pierce BCA Protein Assay Kit with a microplate reader.

The proteins (50 µg) were reduced with dithiothreitol (50 µg) for 1 hour at room temperature (RT), then alkylated with iodoacetamide (125 µg) for 30 minutes in the dark. Reduced and alkylated proteins were bound to Sera-Mag magnetic carboxylate beads (100 µg each), mixed with ethanol, and incubated at RT. Beads were separated magnetically, washed five times with 80% ethanol, and transferred to new tubes. Proteins were digested on-beads with Pierce Trypsin Protease (1 µg) in an ammonium bicarbonate buffer for 18 hours at 37°C while shaking at 1000 rpm. Peptides were recovered through magnetic separation, washed twice with ammonium bicarbonate buffer, and dried using a SpeedVac. The dried peptides were reconstituted in 2% acetonitrile with 5% formic acid, mixed at 300 rpm for 20 minutes, and 10 µL (containing 10 µg protein) was injected for LC-MS/MS analysis^27^.

Proteomics analysis was performed using LC-MS on a Vanquish Flex UPLC system coupled to an Orbitrap Q Exactive Classic mass spectrometer. Peptides were separated on an Agilent AdvanceBio Peptide Map column using a gradient of 2% to 45% buffer B (0.1% formic acid in acetonitrile) over 80 minutes at a flow rate of 0.3 mL/min. Data was acquired in Full MS–SIM and DIA modes.

Raw proteomics data were analyzed with DIA-NN (v1.8.2) in library-free mode using the UniProt mouse reference proteome (February 2024 update)^28^. Cysteine residues were treated as fixed modifications, while methionine oxidation and N-terminal acetylation were variable. Peptide matching was performed with a 1% false detection rate, requiring at least one proteotypic peptide of 7 to 30 amino acids. MaxLFQ intensities were analyzed using Perseus. Protein groups were log-transformed and categorized based on data completeness.

For complete datasets, comparisons were made using a linear model (LIMMA) with false discovery rate correction (q < 0.05). For missing values datasets, Student’s t-test was applied (p < 0.01). Data visualization was performed in R using dplyr, ggplot2, and heatmaps. Functional analysis included pathway enrichment via Metascape (Reactome) and SRPLOT^29^.

#### 2.2.10 Procedure and results of immunization of mice with VLPSi-OVA

Female BALB/c mice, aged 6 to 8 weeks, were randomly assigned to six groups (n = 5 per group). The groups included PBS and OVA immunization as negative controls, alum-OVA as the positive control, VLPSi-5-OVA, and VLPSi-30-OVA as nano-vaccine groups. Subcutaneous immunization was administered with an OVA concentration of 25 µg/mouse where it applied. To enhance the immune response, all groups received booster doses with the same formulation and dosage at 14 and 28 days post-initial immunization. Mouse body weights were recorded every two days following immunization.

#### 2.2.11 ELISA for anti-OVA-IgG

Serum samples were collected at predetermined intervals by obtaining blood from the tail vein, which was subsequently stored at 4 °C overnight. The samples were centrifuged at 3000 rpm for 10 minutes to collect the serum. Remove the ELISA plate from the Chondrex kit and dispense 100 µl of OVA solution into each well. Incubate the plate overnight at 4 °C. Subsequently, wash the plate at least three times with 1 X wash buffer. Introduce 100 µl of Blocking Buffer into each well and incubate for 1 hour at room temperature. Again, wash the plate at least three times with 1 X Wash Buffer. Prepare the standard and sample dilutions, add them to the respective wells, and incubate for 2 hours at room temperature. Following this, add 100 µl of secondary antibody solution to each well and incubate for 1 hour at room temperature. Wash the plate at least three times with 1 X Wash Buffer. Add the TMB solution and incubate for 25 minutes at room temperature. To terminate the reaction, add 50 µl of 2N sulfuric acid (stop solution) to each well. Measure the optical density at 450 nm.

#### 2.2.12 Expression and purification of rEsxB antigen

The antigen rEsxB was chosen as a model to investigate the adjuvant properties of nanoparticle vectors. His-tagged rEsxB was prepared using the standard IPTG induction protocol^30^. In brief, the full-length wild-type EsxB DNA was subcloned into the pET-28a vector and expressed as a 6× His-tag fusion protein in BL21 (DE3) S. aureus. The expression of rEsxB was induced by IPTG and subsequently purified from bacterial cell lysates using Ni-Sepharose chromatography. Ni-Sepharose was eluted using a gradient of imidazole concentrations at 100, 300, and 500 mmol/L separately. Subsequently, the samples underwent dialysis against PBS, and endotoxin removal was performed using a high-capacity endotoxin removal column. The rEsxB solution was then filtered through a 0.22 µm sterile filter, and the total protein concentration was determined via the BCA protein assay. The purity and specificity of the rEsxB antigen were assessed using SDS-PAGE.

#### 2.2.13 Determination of protein loading and cellular safety of NPs-rEsxB

The rEsxB was chemically conjugated through a condensation reaction between the amino groups on the NPs and the carboxyl groups of the rEsxB. For this purpose, 1 mg of rEsxB protein was dissolved in 10 mL of 25 mmol/L MES buffer, followed by the addition of 0.4 mL of 1 mol/L EDC and 0.25 mL of 1 mol/L NHS. The reaction was allowed to proceed for 4 hours at room temperature to activate the carboxyl groups on the rEsxB protein. Subsequently, the NPs were centrifuged at 16,500 × g at 4°C for 1 hour and washed three times with deionized water. The activated rEsxB protein was resuspended in 1 mL of 10 mg/mL NPs in PBS (pH 8.0) and incubated at 4°C overnight. Finally, the NPs-rEsxB conjugates were centrifuged at 16,500 × g at 4°C for 1 hour, washed three times with deionized water, and stored at 4°C for future use. The total loading concentration of rEsxB was quantified using the BCA protein quantitation method. The loading efficiency was determined using the formula: Loading efficiency initial (%) = ((Loaded Mass of rEsxB (mg) / Mass of input rEsxB) × 100%.

BMDC cells were cultured in 96-well plates at a density of 10,000 cells per well. A blank control was established, and two types of nanoparticles PEG-VLPSi-NH_2_-rEsxB-5 and PEG-VLPSi-NH_2_-rEsxB-30 (abbreviated as VLPSi-5-rEsxB and VLPSi-30-rEsxB) were prepared in four experimental groups with concentrations of 0.05 mg/mL, 0.025 mg/mL, 0.0125 mg/mL, and 0.00625 mg/mL, each with five replicate wells. In each group, 100 µL of nanoparticles at varying concentrations were added and incubated for 24 hours. Subsequently, CCK8 reagent was added to achieve a final concentration of 10%, and the OD at 450 nm was measured after a 2-hour incubation period, protected from light. Cell viability was calculated using the formula: Cell Viability (%) = [(OD_test_ – OD_background_) / (OD_control_ – OD_background_)] × 100%.

#### 2.2.14 Effect of NPs-rEsxB on BMDC maturation

BMDCs were cultured in 48-well plates. Experimental groups included a PBS group, an rEsxB group, a VLPSi-5-rEsxB group, and a VLPSi-30-rEsxB group. According to the group assignments, 500 μL of samples were added, and the cultures were incubated for a total duration of 18 hours.

After the incubation period, the cell precipitates were collected and resuspended in 0.4 mL of RPMI 1640 medium. Subsequently, CD40, CD80, and CD86 were added, and the mixture was incubated for 20 minutes at 4°C, shielded from light. Upon completion of the incubation, an additional 0.5 mL of RPMI 1640 medium was introduced, followed by centrifugation at 300 g for 5 minutes. The supernatant was discarded, and this process was repeated three times. A volume of 0.5 mL of fixative was then added, and the samples were analyzed using flow cytometry (NovoCyteTM and Agilent).

#### 2.2.15 NPs-rEsxB nano-vaccine immunized mice

Female BALB/c mice, aged 6 to 8 weeks, were randomly assigned to five groups (n = 15 per group) for immunization. The negative control group received PBS and rEsxB, while the positive control group received Alum-rEsxB. The nano-vaccine groups received VLPSi-5-rEsxB and VLPSi-30-rEsxB separately. Subcutaneous immunizations were conducted with a consistent rEsxB concentration of 50 µg per dose, except for the PBS group. To augment the immune response, all experimental groups received a booster dose of the identical formulation, comprising an rEsxB concentration of 25 µg per mouse, administered at 14 and 28 days subsequent to the initial immunization. Serum samples from the mice were collected at predetermined intervals. To enhance the immune response, all groups were boosted with the same formulation (rEsxB concentration of 25 µg/mouse) at 14 and 28 days after the first immunization. Mouse serum samples were collected at specific time points.

#### 2.2.16 ELISA to measure antibody titer

The rEsxB-specific antibody titer was quantified using an indirect ELISA. In brief, 96-well ELISA plates were coated with rEsxB at a concentration of 500 ng per well and subsequently blocked with 3% BSA the following day. The plates underwent five washes with PBS containing 0.1% Tween 20 (PBST) before being incubated with serially diluted serum from immunized mice (ranging from 1:200 to 1:20,4800; 100 μL per well) for 1 hour at 37°C. Following this incubation, the plates were washed five times with PBST. Horseradish peroxidase (HRP)-conjugated goat anti-mouse IgG (diluted 1:50,000; 100 µL per well) was then added and incubated at 37 °C for 45 minutes, followed by a washing step as previously described. ABTS and hydrogen peroxide (H_2_O_2_) served as substrate chromogens, and color development was allowed to proceed in the dark for 30 minutes at room temperature. Absorbance measurements were conducted at a wavelength of 405 nm using a microplate reader (Biotek SYNERGY LX Instruments, Santa Clara, CA, USA). The cut-off value was determined as 2.1 times the absorbance of the negative control serum, as referenced in^31^. In comparative analyses of ELISA titer results among sample groups, the efficacy of the area under the curve (AUC) method was comparable to that of the absorbance summation method and surpassed that of the end-point titer method. The rEsxB-specific antibody titer results were reported as the mean serum ELISA AUC values^32^, with each group consisting of 15 mice.

#### 2.2.17 ELISA for measurement of IL-4 and INF-γ levels in mouse serum

Mouse serum was collected consistently as described in Method 2.2.11. A 1.5 μL aliquot of each mouse serum was diluted 100-fold to a final volume of 150 μL using a sample diluent. Various concentrations of Mouse INF-γ standards (500, 250, 125, 62.5, 31.25, 15.6, 7.8, 0 pg/mL) were prepared. Separate wells were designated for zero controls, labeling, and sample diluent testing. The plate was covered with a sealing film and incubated at 37°C for 2 hours. Subsequently, the plate was washed 3-4 times with 350 μL per well of 1× Wash Buffer and dried. Each well received 100 μL of 1× Anti-Mouse IFN-γ Detection Antibody and was incubated at 37 °C for 1 hour, followed by another round of washing 3-4 times. Then, 100 μL of HRP-conjugated antibody was added to each well and incubated at 37 °C for 40 minutes, with washing repeated 3-4 times. Finally, 100 μL of TMB substrate was added to each well, and color development was conducted at 37 °C for 15-20 minutes, protected from light. A volume of 100 µL of Stop solution was added to each well, and the OD value of each well was measured at 450 nm using an ELISA reader. The readings were recorded within five minutes following the addition of the Stop solution. The level of interleukin-4 (IL-4) cytokine was quantified using the aforementioned method.

#### 2.2.18 ELISpot measures the number of splenocytes expressing IL-4, IFN-γ

On day 7 post the third immunization, spleens were collected from mice of five groups and processed into single-cell suspensions. ELISPOT plates, pre-coated with antibodies, were washed with PBS and incubated with RPMI-1640 complete medium (10% FBS) for 30 minutes at room temperature. Splenocytes (2.5 × 10^6^ cells/well) were mixed with 4 µg/mL recombinant EsxB and added to the plates, followed by incubation at 37 °C for 24 hours. PHA (10 µg/mL) and medium served as positive and negative controls. After incubation, cells were removed, and plates were washed. Detection antibody in PBS with 0.5% FBS was added for 2 hours at room temperature. Following washes, TMB substrate was added until spots appeared. The reaction was stopped by washing with deionized water, and plates were dried in the dark. Spots were quantified using an ELISPOT reader (Cellular Technology Limited, USA).

#### 2.2.19 Lethal Challenge Experiment Evaluates the Ability of NPs-rEsxB Nano-vaccine to Prevent Staphylococcus aureus Infections

On day 7 post the third immunization, each group of mice (n=10) was intravenously injected with 100 µL of Staphylococcus aureus (ATCC 25923) at 6.4 × 10^8^ CFU/mL. Mice were monitored daily for mortality, vital signs, and overall health status over a two-week period. Surviving mice were then euthanized, and body weights were recorded. Viscera were collected for hematoxylin and eosin (HE) staining analysis. In the second lethal challenge, the intravenous dose of Staphylococcus aureus was 12.8 × 10⁸ CFU/mL.

### 2.3. Statistical Analysis

All statistical analyses were studied by using GraphPad Prism 10.3.0 software. Results were expressed as means ± SD. Differences between two groups were tested using an unpaired, two-sided Student’s T test. Differences among more than two groups were evaluated by one-way ANOVA with significance determined by Tukey-adjusted t-tests.

## 3. Results

### 3.1 Physicochemical characteristics, antigen release, and enhanced cell uptake of VLPSi

TEM images show the both VLPSi-5 and VLPSi-30 have virus-like shape and possess 5 and 30 nm surface spikes in length. The reference sample VLPSi-0 is spherical with smooth surface (Figure 1a-c). The plain VLPSi show negative surface charge with similar value between –23 and –28 mV regardless of surface spike length (Figure 1d). After PEGylation, the zeta potential of the VLPSi increased by approx. 2- 4 mV. Amino groups modification on the VLPSi led to a charge reversal from negative to positive. The surface charge displayed + 35.3, + 34.2, + 31.5 mV for PEG-VLPSi-NH_2_-0, PEG-VLPSi-NH_2_-5, PEG-VLPSi-NH_2_-30, respectively. Hence, zeta potential measurements confirmed the successful surface functionalization on the VLPSi. The OVA loadings for VLPSi-30-OVA and VLPSi-5-OVA were determined to be 8.25% and 7.85% (Table S1, supporting information). The OVA release from different VLPSi was studied at 37 °C for 24 h *in vitro* (Figure 1e). The OVA was released from the VLPSi in a sustained manner and the length of the surface spikes did not significantly affect the release profiles of OVA.

**Figure 1.**
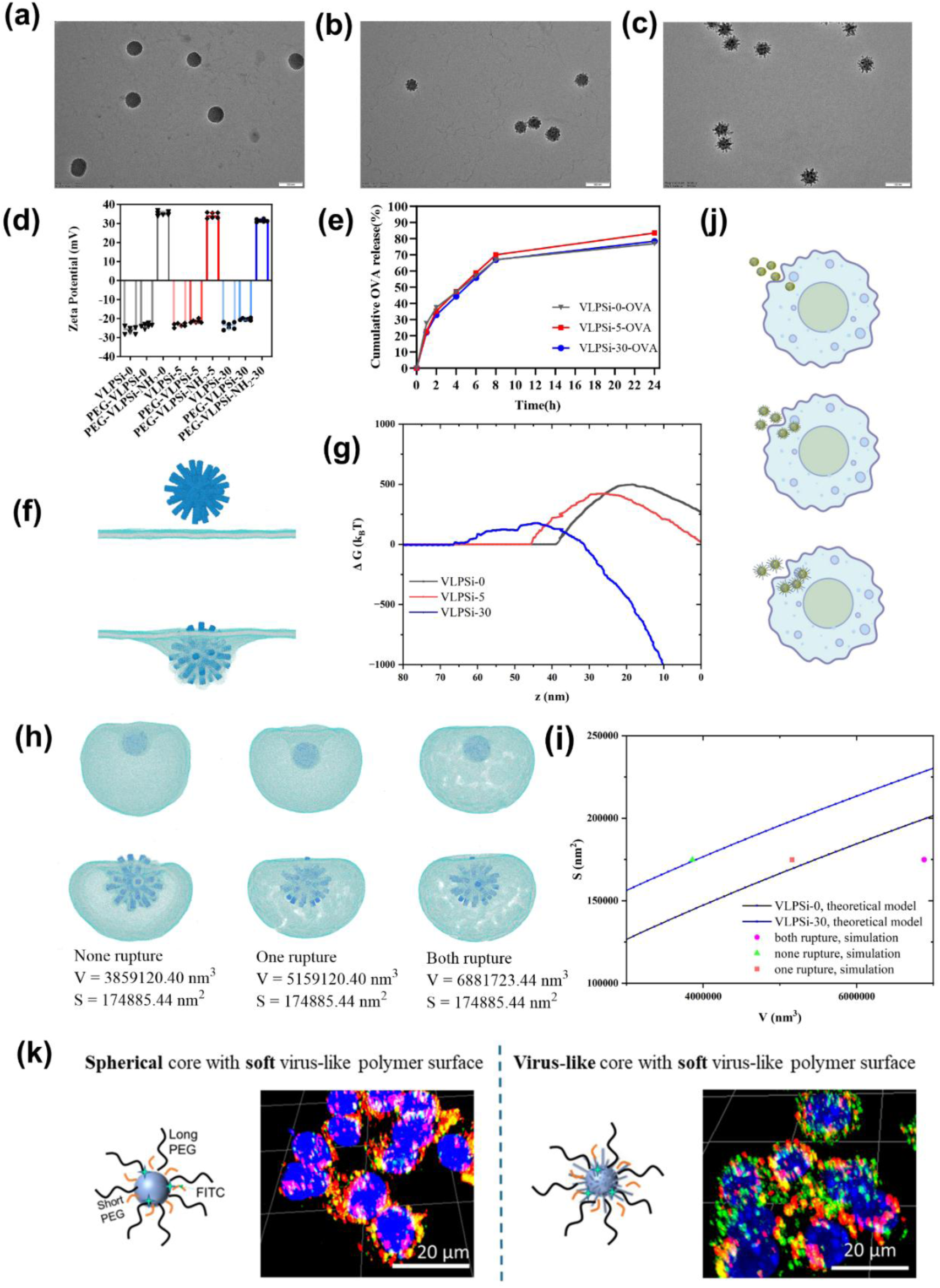
Synthesis and structural characterization of virus-like mesoporous silica nanoparticles. (a)TEM images of VLPSi-0. (b) VLPSi-5. (c) VLPSi-30. The scale bars are 100 nm. (d) Zeta potential analysis VLPSi with different spike length and surface modifications. (e) The release percent of OVA from VLPSi with varying spike lengths in a PBS solution at pH 7. The data are presented as mean ± standard deviation (SD), n=3. (f) Typical CGMD simulation snapshot of spiky NP across the cell membrane. (g) Change of free energy across membrane of NPs with different spiky lengths. (h) Typical CGMD simulation snapshots of the rupture of lysosomes encapsulating of spiky NP. (i) Theoretical prediction and simulation results of lysosomal engulfment of NP. (j) Schematic illustration for cellular uptake of three types of VLPSi. (k) Confocal microscopy images for studying cell uptake and endosomal escape of VLPSi-0 and VLPSi-30 uptake with soft spiky PEG coating. The VLPSi were labeled with FITC and incubated with RAW264.7 for 4 hours, with the fluorescence signal shown in green. The nucleus and lysosome were stained with DAPI (blue) and Lysotracker (red). The scale bar represents 20 µm.

The interaction between VLPSi and immune cells, such as macrophage cells, is crucial for antigen presentation to activate immune response post-vaccination. To thermodynamically illustrate the important role of spike in the interaction between NPs and cell membranes, we calculated the changes of free energy of NPs as they cross the cell membranes. As shown in Figures 1f and 1g, the presence of spikes on NPs significantly reduces the free energy barrier for them to cross the cell membranes, with a significant value of ∼319𝑘_B_𝑇 for VLPSi-30 as compared to VLPSi-0. Next, we simulated and compared the lysosomal escape efficiency of spiky NPs versus spherical NPs. Taking the escape of a single NP in the lysosome as an example, we start with a lysosome capable of completely encapsulating the NP with 𝑆 = 1.75 × 10^6^ nm^2^ and 𝑉 = 3.86 × 10^7^ nm^3^. As the lysosomal volume increases to 5.16 × 10^7^ nm^3^, the lysosomes that engulf the spiky NPs will rupture, while the lysosomes that engulf the spherical particles remain intact. And the lysosomal rupture happens for both conditions when the volume increases to 𝑉 = 6.88 × 10^7^ nm^3^. As shown in Figure 1h, we compare the simulation results and the theoretical predictions. The solid lines in Figure 1i represent the limit size of the lysosome that can engulf a given NP predicted by the Eq. (4). The area above the solid line indicates the size of the lysosomes that can engulf a NP and maintain their integrity, while the area below the solid line indicates that it cannot. Based on the above simulation results, it can be inferred that spiky VLPSi are more likely to escape lysosomes than spherical NPs.

To verify the simulation results, we conducted cell internalization experiments by incubating FITC-labeled NPs with RAW264.7 cells. The order of internalized VLPSi into RAW264.7 cells was VLPSi-30 > VLPSi-5 > VLPSi-0, indicating that longer surface spikes on the VLPSi could enhance cellular uptake (Figure S1, supporting information). To study the effect of spike stiffness on cell uptake and endosomal escape, a spiky PEG coating was functionalized on VLPSi by using two different lengths of PEG-silanes (0.5 and 2.0 kDa). The organic PEG spikes are soft while inorganic SiO_2_ spikes on VLPSi are rigid. As shown in Figure 1k, the spherical VLPSi-0 with soft spiky PEG coating has much less cellular uptake than the VLPSi-30 with the same coating. The confocal images of VLPSi-0 present many bright yellow dots which are due to overlapping green FITC labelling and red Lysotracker staining. This indicates that the internalized spherical VLPSi-0 NPs were mainly trapped in endosome/lysosome. In contrast, VLPSi-30 NPs effectively escape from endosomal after internalization and thus many green dots are observed in the 3D confocal images. These findings demonstrate that rigid spikes on NPs can significantly promote both cellular uptake and endosomal escape, while soft rigid spikes fail to produce the same effect.

### 3.2 Immunoadjuvant properties *in vitro*

Macrophages are one of the most important and first-interacted immune cells when NPs are administrated. Thus, the interaction mechanism between the OVA loaded VLPSi with macrophages was studied with proteomics. Figure 2a shows the schematic proteomics of the three VLPSi samples. In the volcano of differentially expressed proteins, the groups exposed to the VLPSi-0-OVA, VLPSi-5-OVA, and VLPSi-30-OVA have 293 (121 up-regulated and 172 down-regulated proteins), 269 (117 up-regulated and 152 down-regulated), and 254 (115 up-regulated and 139 down-regulated) differentially regulated proteins, respectively (Figure 2b-d).

**Figure 2.**
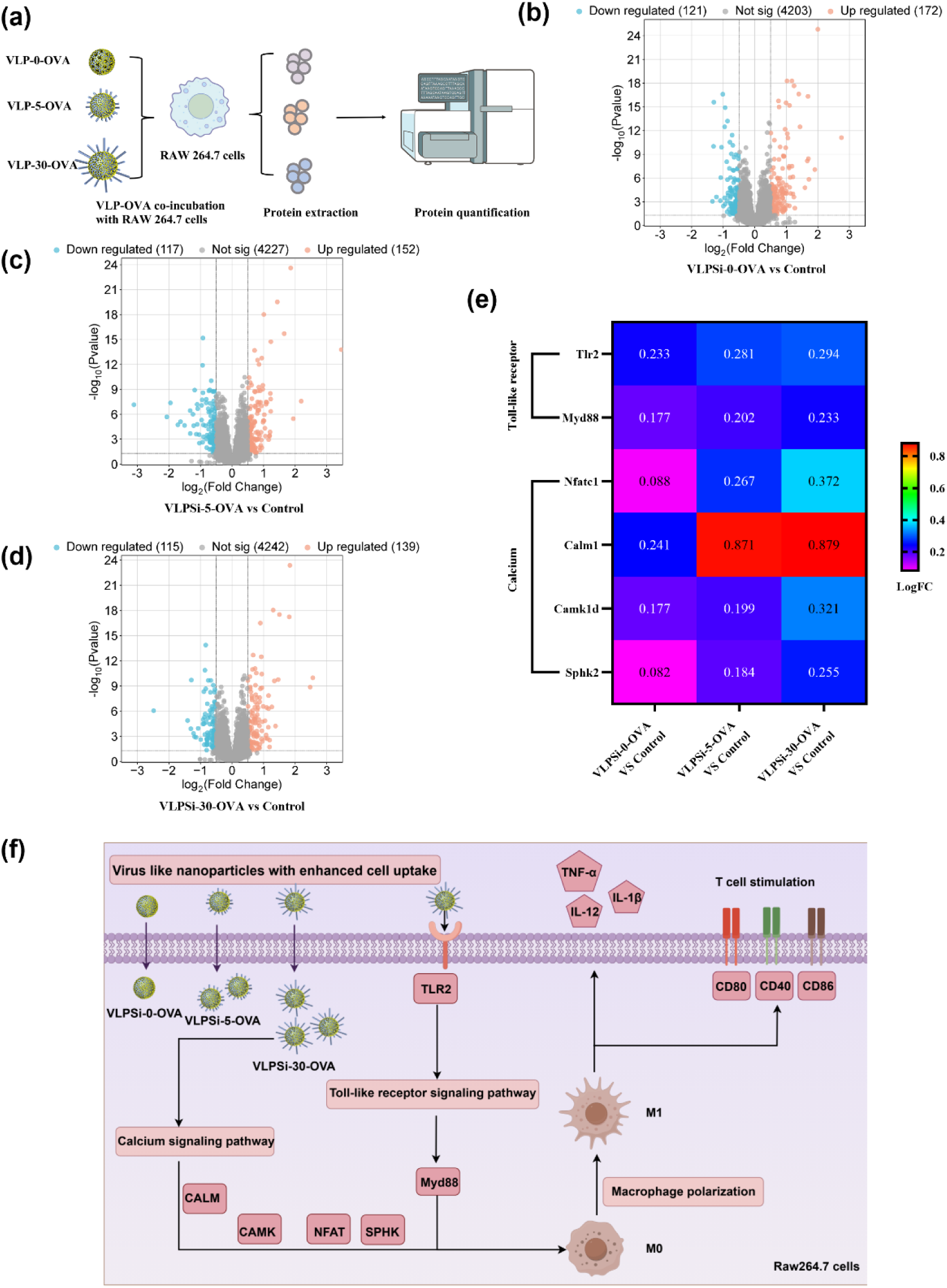
(a) Schematic illustration for proteomics of three types of VLPSi. (b, c, d) Volcano plot of differentially expressed proteins in RAW 264.7 after incubated with VLPSi-OVA with spike lengths of 0, 5, and 30 nm. The red dots represent the upregulated proteins, the blue dots represent the downregulated proteins, and the gray dots represent no significant change (P < 0.05, |log_2_FC|>0.5). (e) Proteins upregulated in the calcium and Toll-like receptor signaling pathways in the VLPSi-0-OVA, VLPSi-5-OVA, and VLPSi-30-OVA groups relative to the control group, with more pronounced protein upregulation corresponding to longer surface spikes. (f) Schematic illustrates the specific locations and functions of these upregulated proteins within the calcium signaling pathway and TLR signaling pathway.

Pathway enrichment analysis of sample-treated macrophages is shown in Figure S2 (Supporting information). The results indicate that VLPSi-0-OVA mainly activated core innate immune mechanisms (Figure S2a), including inflammasomes, NF-κB, interleukin 1 signaling, and TLR pathways. VLPSi-5-OVA induced broader immune activation than VLPSi-0-OVA. In addition to innate pathways, macrophages showed enrichment of MHC class II antigen presentation and CaMK signaling pathways, indicating improved antigen uptake and adaptive immune activation^33^. VLPSi-30-OVA generated the most extensive activation, with strong enrichment of innate immune signaling, antigen presentation, and lymphoid–myeloid communication pathways. Calcium dependent signaling trafficking and Rho GTPases pathways were more prominent than in VLPSi-5-OVA. Additional enrichment of extracellular matrix remodeling further indicated a highly activated macrophage state. Overall, these results support increasing spike length produced a clear enhancement of immune activation.

Figure 2e presents a heatmap illustrating the logFC values for VLPSi-0-OVA, VLPSi-5-OVA, and VLPSi-30-OVA, compared to the protein expression levels in the control group. The results suggest that the expression levels of proteins such as Tlr2 and Myd88 within the TLRs signaling pathway, alongside Nfatc1, Calm1, Camk1d, and Sphk2 within the calcium signaling pathway, exhibit a positive correlation with the length of VLPSi surface spikes. Specifically, elongated VLPSi surface spikes are associated with enhanced expression of these proteins. Figure 2f illustrates these upregulated proteins within the TLRs and calcium signaling pathways. This enhancement suggests that the VLPSi with long spikes has the potential to facilitate the increased secretion of pro-inflammatory molecules (e.g., IL-1β and TNF-α) and promoted T cell activation through the upregulation of these proteins.

The efficacy of OVA-loaded VLPSi in the activation of BMDCs is shown in Figure 3. Figure 3a shows a schematic diagram of the effects of the three VLPSis on BMDC maturation. At first, none of OVA loaded NPs exhibited cytotoxicity towards BMDC cells (Figure 3b-d) or HUVEC cells in the concentration of 25-200 µg/mL (Figure S3a-c), as compared to the controls. Thus, the NPs demonstrated good biocompatibility *in vitro*. Then, BMDCs were incubated with different samples for 24 hours and then the surface expression of CD40 (Figure 3e), CD80 (Figure 3f), and CD86 (Figure 3g) on BMDCs were quantified using flow cytometry (Figure S4). The findings indicated that BMDCs treated with VLPSi-30-OVA exhibited higher degree of maturation than those treated with the VLPSi-5-OVA. The sample VLPSi-0-OVA group showing the least maturation (Figure 3e-3g). Additionally, the concentrations of cytokines, including IL-6 (Figure 3h), IFN-γ (Figure 3i), and IL-1β (Figure 3j) in the supernatants of BMDCs from each group were determined using ELISA. The group of VLPSi-30-OVA presented the highest IFN-γ levels among the tested groups, followed by VLPSi-5-OVA group (VLPSi-5-OVA vs. VLPSi-30-OVA p=0.0009). The VLPSi-0-OVA group shows the lowest ability to evoke the expression of IFN-γ (VLPSi-0-OVA vs. VLPSi-5-OVA p<0.0001). Regarding IL-1β, both VLPSi-30-OVA and VLPSi-5-OVA promoted similar expression level, and both had higher IL-1β concentrations

**Figure 3.**
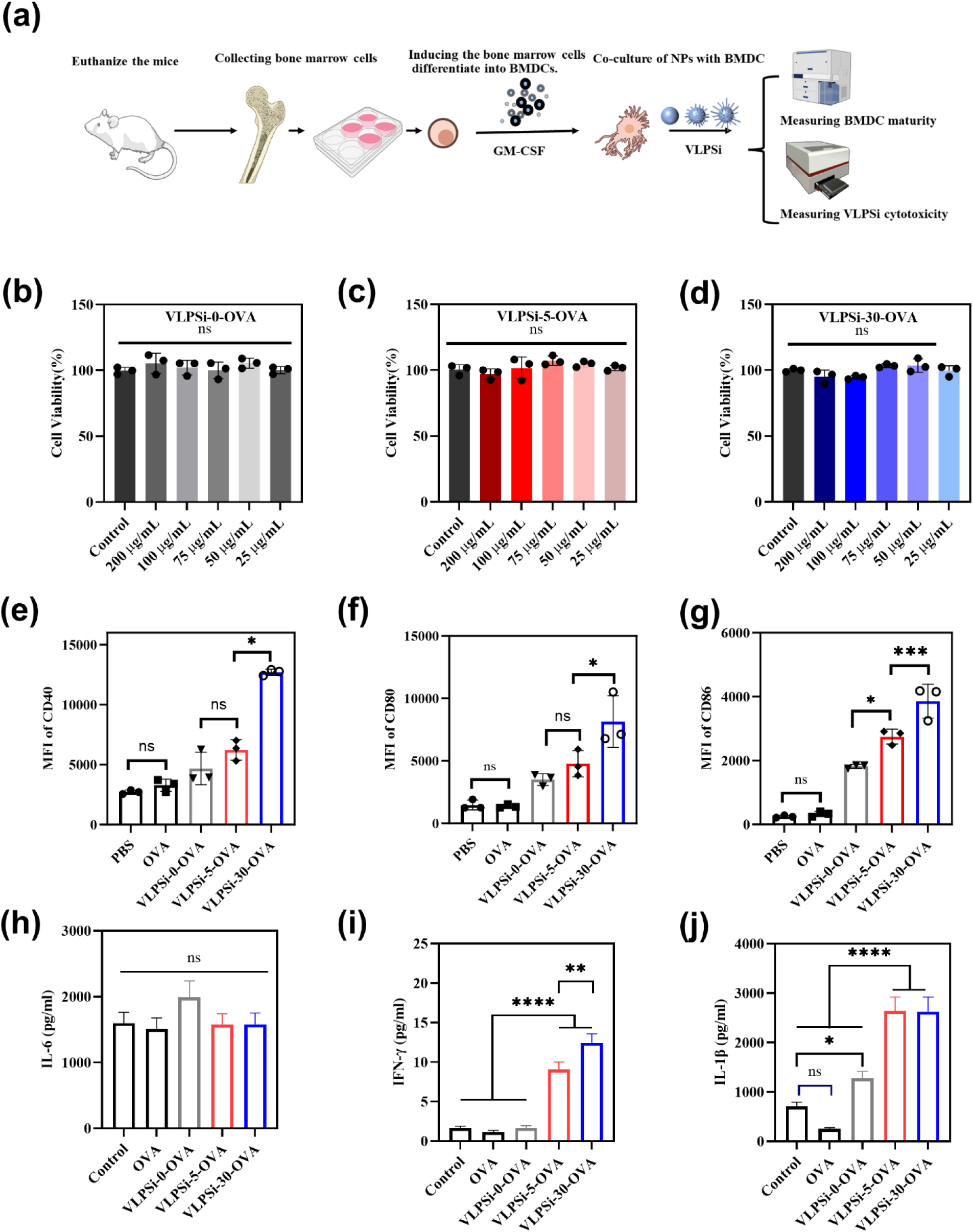
(a) Schematic illustration for BMDCs collection and three types of VLPSi to induce BMDC maturation, cytotoxicity and cytokine secretion. (b, c, d) The toxicity of BMDCs induced by VLPSi-0-OVA, VLPSi-5-OVA, and VLPSi-30-OVA. (e, f, g) The impact of VLPSi-0-OVA, VLPSi-5-OVA, and VLPSi-30-OVA on the maturation of BMDCs. (h, i, j) The effects of OVA-loaded VLPSi on IFN-γ, IL-1β, IL-6 secretion by BMDCs (* *p* < 0.05; ** *p* < 0.01; *** *p* < 0.001; **** *p* < 0.0001) than the VLPSi-0-OVA group (p <0.0001). No statistically significant differences were observed in IL-6 levels across the groups (p>0.05). The increased expression of IL-6 is generally related to side effects of vaccines^34^. Thus, these results indicated that OVA-loaded NPs had good safety features although they are effective in elevating the proinflammatory cytokines such as IFN-γ and IL-1β for immune activation.

### 3.4 Immunoadjuvant properties of nanomaterials in vivo

VLPSi-30-OVA and VLPSi-5-OVA were selected to assess their immune activation potential *in vivo*. The experimental procedure is described in Figure 4a. VLPSi-30-OVA exhibited a significantly greater capacity to enhance the expression of Anti-OVA IgG compared to VLPSi-5-OVA (Figure 4b), with a p-value of 0.0435. No statistically significant difference was observed in IgG production between VLPSi-5-OVA and aluminum (p = 0.9882). Following the ELISA analysis, it was observed that VLPSi-30-OVA and VLPSi-5-OVA elicited significantly higher levels of IL-2 in mice compared to the aluminum group (Figure 4 c), potentially attributable to the activation of cellular immunity by VLPSi nanomaterials (Alum-OVA vs. VLPSi-5-OVA, p = 0.0010). However, the aluminum group did not exhibit a significant increase in IL-2 levels compared to the control group (p = 0.7679). In the Elispot assay (Figure 4 d and Figure S5), both VLPSi-30-OVA and VLPSi-5-OVA significantly elevated the number of IL-4 secreting splenocytes (Alum-OVA vs. VLPSi-5-OVA, p = 0.0011), with no significant difference observed between VLPSi-30-OVA and VLPSi-5-OVA (p = 0.1819). The aluminum group also showed an increase in IL-4-secreting splenocytes compared to the control and OVA groups (p < 0.0001). Regarding IFN-γ secreting splenocytes (Figure 4e and Figure S6), VLPSi-30-OVA demonstrated a stronger induction capacity than VLPSi-5-OVA (p = 0.0312), while no significant increase was noted in the aluminum group relative to the control group (p = 0.4901). Both groups exhibited good biosafety, as indicated by the lack of significant differences in body weight and vital signs among the mice in each group (Figure S7).

**Figure 4.**
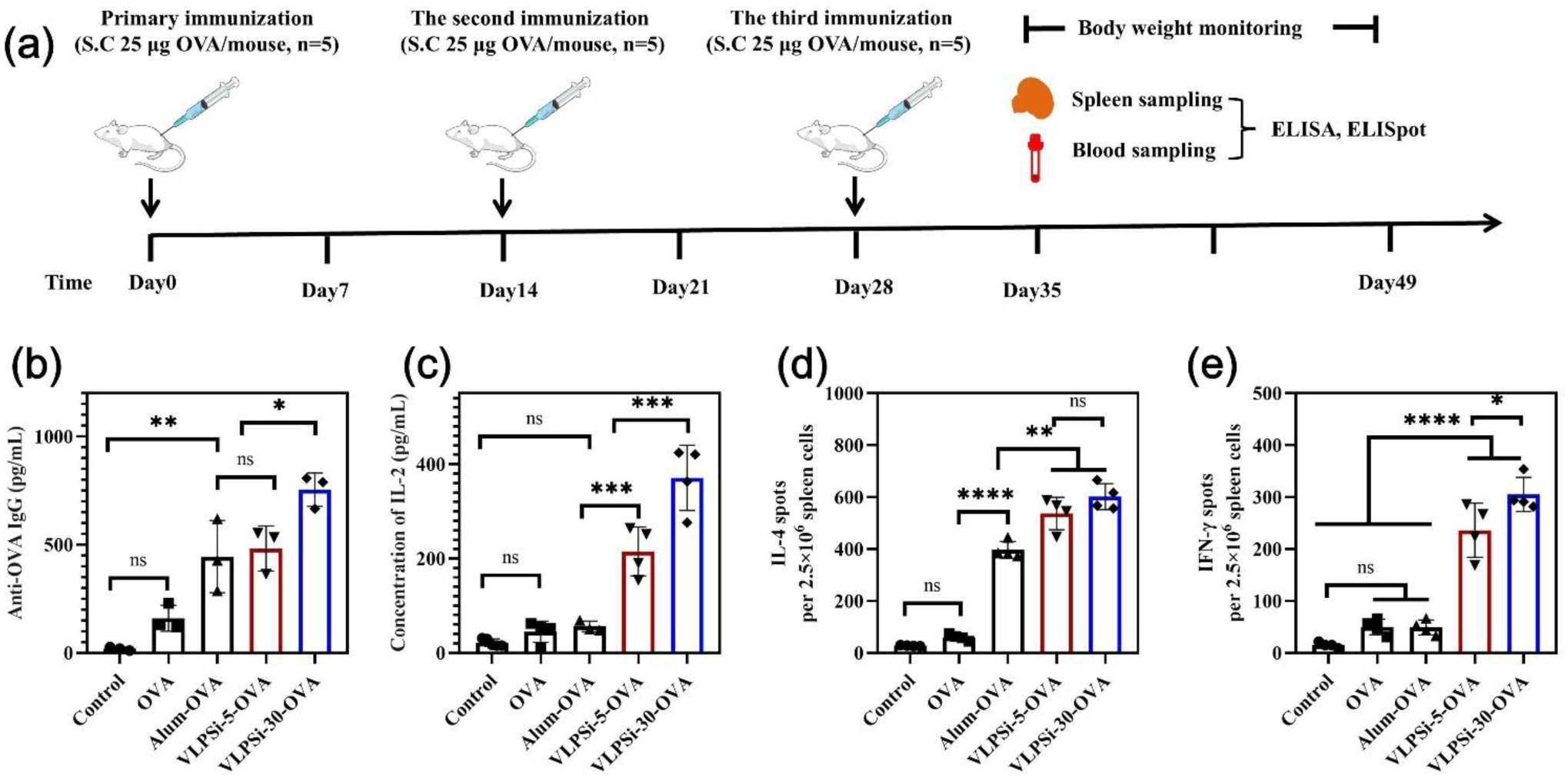
(a) The flow chart of NPs-OVA immunized animals. (b) The serum levels of Anti-OVA IgG as measured by ELISA. (c) The serum levels of IL-2 measured by ELISA. (d, e) The ELISpot assay results for the secretion of IL-4 and IFN-γ by splenic cells. The P-value of less than 0.05 was considered statistically significant (* *p* < 0.05; ** *p* < 0.01; *** *p* < 0.001; **** *p* < 0.0001).

### 3.5 VLPSi-based nanovaccines against bacterial infection

These preliminary experiments demonstrated that VLPSi, when loaded with the OVA antigen, could induce a stronger immune response than Alum. Subsequently, we assessed the antimicrobial efficacy of the VLPSi vaccine in comparison to the Alum vaccine by loading the recombinant EsxB (rEsxB) antigen. In the following study, the antigen of staphylococcus aureus and rEsxB was covalently bonded on VLPSi via amide linkages. The loading degree of rEsxB on VLPSi is shown in Table S2. *In vitro* cellular assays further verified that VLPSi-rEsxB was not cytotoxic (Figure S8). Then, the expression of CD40, CD80, and CD86 on the surface of BMDCs was determined by flow cytometry after co-incubation of VLPSi-rEsxB with BMDCs. This experiment confirmed that MSN induced a stronger immune response compared to aluminum. As demonstrated in Figures S9 a-d, the flow cytometry assays revealed that the “spikes” on the NPs’ surface significantly enhanced BMDC maturation. This result is consistent with the results of VLPSi-OVA. The results indicated that VLPSi loaded with the rEsxB antigen confirmed the hypothesis that the length of the “spikes” on the surface of nanoparticles is positively correlated with the strength of immune activation.

To further assess the efficacy of nano-adjuvants in preventing pathogenic infections, we conducted a lethal challenge experiment. This setup was employed to evaluate the protective capabilities of two nanomaterials, VLPSi-30-rEsxB and VLPSi-5-rEsxB, against mice infection. The experimental procedure for the lethal challenge is depicted in Figure 5a. Both nano-vaccines with VLPSi elicited higher IgG level than those with aluminum. Furthermore, the mean IgG antibody titer induced by VLPSi-30-rEsxB was higher than VLPSi-5-rEsxB although no significant difference was observed between them (Figure 5b). Both VLPSi-30-rEsxB and VLPSi-5-rEsxB elicited significantly higher IgG levels than aluminum (p < 0.001). These results indicated that VLPSi was effective vaccine adjuvant to evoke potent memory immune response and the length of spikes on VLPSi had positive correlation with activated immune response. When studying cytokine expression specifically, VLPSi-30-rEsxB demonstrated superior performance to promote the production of IL-4 (Figure 5c) and IFN-γ (Figure 5d) compared to VLPSi-5-rEsxB. Aluminum-rEsxB presented the weakest results among the tested vaccines. These results were further validated with Elispot assays further corroborated these findings (Figure 5e and Figure 5f), indicating showing that VLPSi-based nanovaccines with long spikes are the best candidate to activate humoral immune response regarding. Both formulations demonstrated superior efficacy compared to Aluminum-rEsxB.

**Figure 5.**
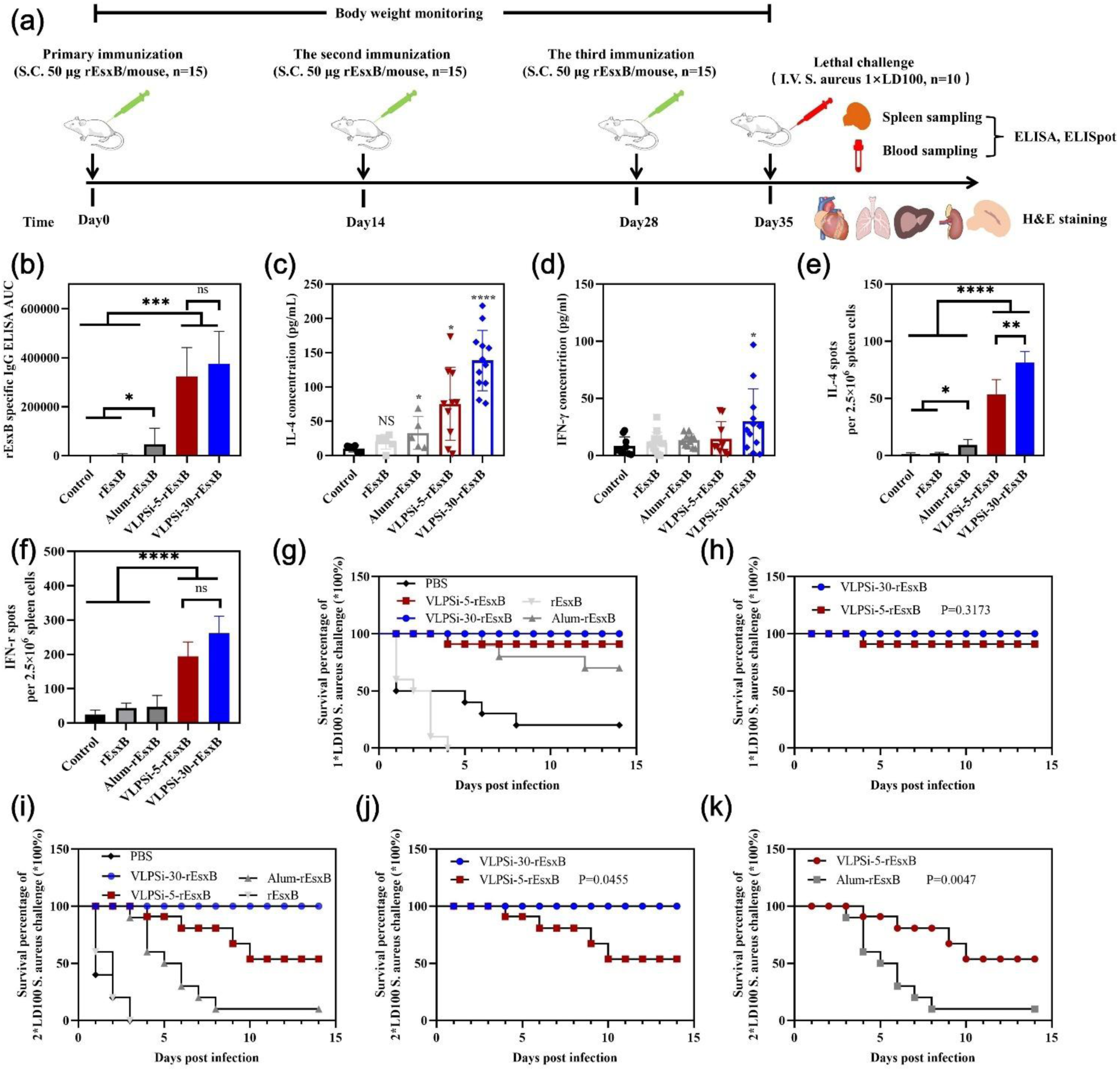
Results of subcutaneous injection of NPs-rEsxB in mice and subjected to lethal challenge with S. aureus. (a) The experimental flow chart for the lethal challenge. (b) The total serum rEsxB-specific IgG titer assay in NPs-rEsxB-immunized mice. (c, d) ELISA quantification of IL-4 and IFN-γ in mice. (e, f) The ELISpot assay results for the secretion of IL-4 and IFN-γ by splenic cells. (g, h) The survival rate of 1×LD100 S. aureus challenge experiment. (i, j, k) The survival rate of 2×LD100 S. aureus challenge experiment. All data are expressed as mean ± standard deviation (S.D.). The P-value of less than 0.05 was considered statistically significant (* p < 0.05; ** p < 0.01; *** p < 0.001; **** p < 0.0001).

To evaluate the protection effect of the activated immune response, lethality experiments were carried out. The results indicated a hierarchy in survival rates among mice, with VLPSi-30-rEsxB exhibiting the highest survival, followed by VLPSi-5-rEsxB, and finally Aluminum-rEsxB, as shown in Figure 5g. However, there was no statistical significance in the survival of mice between these three groups, as shown in Figure 5h and Figure S10. This observation indicates that the initial lethal dose was insufficient to effectively evaluate the efficacy of the three nano-vaccines. Consequently, we increased the challenge dose to 2 × LD100. The experimental procedure remained consistent with that described in Figure 5a, except that the quantity of S.aureus administered intravenously post-immunization was doubled. As illustrated in Figure 5i, all mice in the pure antigen rEsxB and PBS-treated groups succumbed following the challenge of the 2×LD100 dose of S. aureus. In contrast, the VLPSi-30-rEsxB group exhibited a 100% survival rate, while the VLPSi-5-rEsxB group showed a 60% survival rate, with a statistically significant P-value of 0.0455 between these two groups (Figure 5j). The survival rates in both the VLPSi-30-rEsxB and VLPSi-5-rEsxB groups were significantly higher compared to the alum vaccine group, where only 10% of the mice survived. The survival analysis between the VLPSi-5-rEsxB and Alum-rEsxB groups yielded a P-value of 0.0047 (Figure 5k). Detailed survival outcomes for the remaining groups are provided in Figure S11.

The VLPSi-based nano-vaccines exhibited excellent biocompatibility *in vivo*. As shown in Figures 6a and 6b, there were no significant differences in body weight among the groups of mice, and no apparent lesions were observed in the H&E staining of major organs, including the heart, liver, spleen, kidney, and lung.

**Figure 6.**
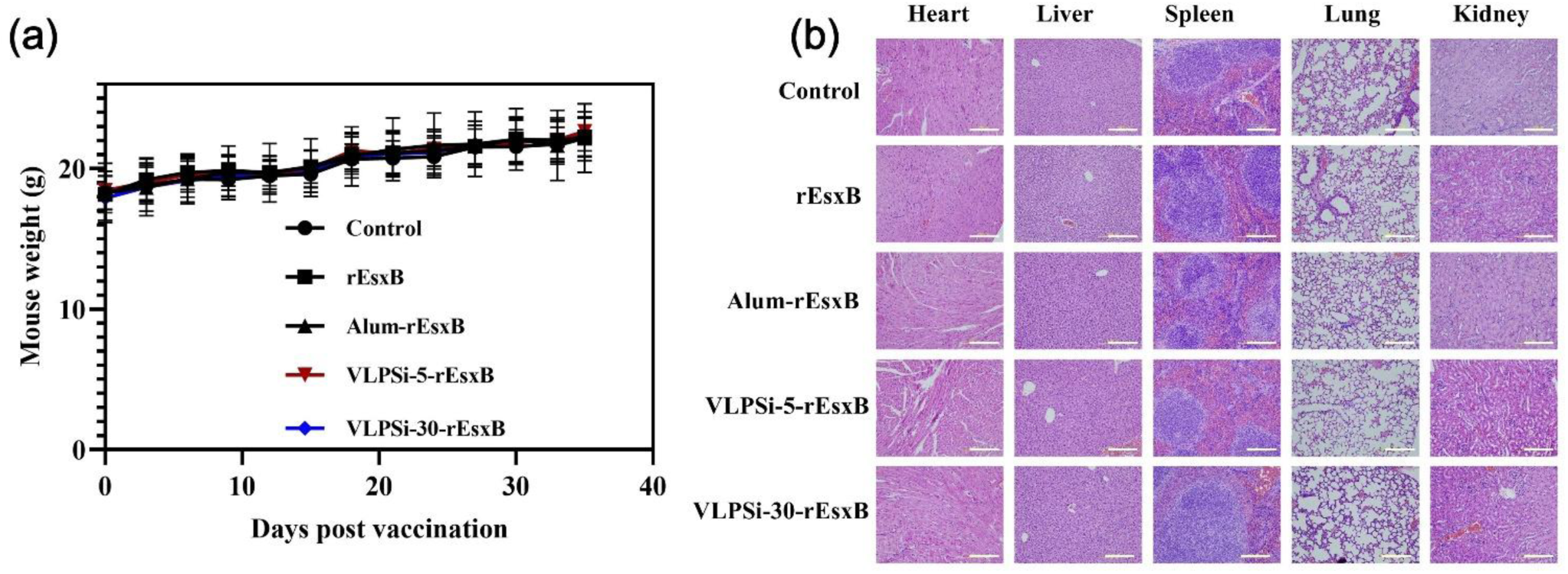
(a) The changes in body weight for each group of mice. Data are represented as means ± SD (n = 10). (b) H&E staining analysis of mouse heart, liver, spleen, lung, and kidney. Scale bars, 200 μm. All data are expressed as mean ± standard deviation (S.D.).

## 4. Discussion

Silicon nanomaterials are gaining attention for their potential as adjuvants in vaccines, enhancing both humoral and cellular immune responses. For instance, they can boost antibody levels and T cell proliferation in vaccines, activating immunity through TLR pathways^35^. Also, silica NPs are being explored for allergen-specific immunotherapy due to their biodegradability and ease of surface modification, which can improve allergy treatment by targeting antigen-presenting cells^36^. In cancer immunotherapy, silicon-based nanomaterials like hollow mesoporous silica nanoparticles can stimulate Th1 antitumor immunity and promote immunological memory, offering a pathway to more effective and targeted cancer immunotherapies^37^. Despite promising progress, utilizing VLPSi as the immune adjuvant and its mechanism in immune activation have been rarely investigated.

In this study, VLPSi NPs, which possess biomimetic morphology of viruses, were studied as novel immune adjuvant. Initially, OVA was used as the model antigen of our investigation. OVA was loaded on the NPs based on electrostatic interaction. Due to the piercing effect and mechanical pressure of the nanospikes, the NPs disrupt the stability of cell membrane. Also, Coarse-grained molecular dynamics simulation demonstrates the presence of spikes on NPs significantly reduces the free energy barrier for them to cross the cell membranes. Thus, the NPs enhance cell internalization and activate immune response, which was validated with RAW 264.7 and DCs. The VLPSi-30-OVA with long spike presented significantly higher cell uptake than VLPSi-5-OVA with short spike or VLPSi-0-OVA with spherical shape. This result is consistent with the findings of Ngoc Man Phan et al.^38^ Then, VLPSi-30-OVA effectively induced DC maturation and increased cytokine secretion, thereby activating both humoral and cellular immunity. The immune activation mechanism was explored with proteomics in detail. Proteomic analysis reveals that the spike length of VLPSi nanoparticles has major effects to influence immune activation in macrophages. Compared to spherical particles, spiked NPs (VLPSi-5/30-OVA) preferentially activate pathways for antigen presentation and, crucially, Ca²⁺ signaling—a key regulator of adaptive immunity. The upregulation of TLR proteins with increasing spike length directly links nanostructure to immune potency. Thus, the superior adjuvant effect of VLPSi-30 is mechanistically explained by its ability to activate Ca²⁺ and endosomal TLR pathways, efficiently bridging innate and adaptive immune responses.

The consistency between the rEsxB and OVA models confirms that the immune-enhancing effect is intrinsic to the VLPSi platform, not antigen-specific. The key mechanistic insight is the positive correlation between the length of the surface nanospikes and the strength of immune activation. This is demonstrated by the enhanced maturation of BMDCs and the elevated levels of antigen-specific IgG, IL-4, and IFN-γ induced by VLPSi-30-rEsxB compared to its short-spiked counterpart, VLPSi-5-rEsxB. The ability to stimulate a balanced Th1/Th2 response is a distinct advantage over the typically Th2-skewed alum adjuvant. The most compelling evidence comes from the lethal challenge experiment. Under a stringent 2×LD100 dose, the hierarchy of vaccine protection was clearly established: VLPSi-30-rEsxB (100% survival) > VLPSi-5-rEsxB (60% survival) > Alum-rEsxB (10% survival). The statistically significant survival benefit of VLPSi-30-rEsxB over both VLPSi-5-rEsxB (p=0.0455) and Alum-rEsxB (p=0.0047) directly translates the observed cellular and humoral immune responses into a critical, life-saving outcome.

## Conclusions

In conclusion, this study systematically demonstrates that VLPSi nanoparticles, particularly the long-spiked VLPSi-30, function as a highly effective and versatile adjuvant platform for vaccine development. The biomimetic viral morphology of VLPSi-30, characterized by its pronounced nanospikes, was identified as a critical factor for its superior performance. Key findings reveal that these nanostructures enhance cellular internalization and activate immune responses because the presence of spikes on NPs significantly reduces the free energy barrier to cross the cell membranes and subsequent promotion of antigen uptake by DCs. Proteomics reveals that the enhanced adjuvant effect of spiked VLPSi-30 stems from its specific activation of Ca²⁺-mediated and endosomal TLR immune pathways. The VLPSi-30 potently activated both humoral and cellular immunity when using OVA and rEsxB as antigens. Consequently, the VLPSi-30-rEsxB nano-vaccine achieved 100% survival in the lethal S. aureus challenge, significantly outperforming the alum-adjuvanted vaccine (10% survival). Therefore, this work provides a solid scientific foundation for applying VLPSi as potent adjuvants in next-generation vaccines against a broad spectrum of infectious diseases.

## Conflict of Interest

No potential conflict of interest was reported by the authors.

## Author Contributions

Conceptualization, W. X., F.L.; The experiments design, W. X., F. L.; Synthesis of nanomaterials, J.W.; Synthesis and characterization of the nanovaccines and animal experiments, C.P.; Cell experiment, J. W., C. P.; Proteomics, A.M.; Computation simulation, S. M. and Q. H. Writing—original draft preparation, C.P., J.W.; Writing—review and editing, V. L., F. L., and W. X.; Supervision, W. X., H.L. and F. L.; Resources, W.X., H.L.. The manuscript was written through contributions of all authors. All authors have read the final version of the manuscript and given approval for publication.

## Funding Sources

This work was carried out with financial support from Research Council of Finland (Grant no. 356056) and Program of China Scholarship Council. SIB Labs, UEF Cell and the Tissue Imaging Unit were acknowledged for technical support. We also acknowledge Biocentre Finland for the use of their LC-MS laboratories.

## Abbreviations

APCs: Antigen-presenting cells
AUC: Area under the curve
BSA: Bovine serum albumin
BMDCs: Bone marrow-derived dendritic cells
CCK-8: Cell Counting Kit-8
DAMPs: damage-associated molecular patterns
DC: Dendritic cell
ELISA: Enzyme-linked immunosorbent assay
FBS: Fetal bovine serum
HUVECs: Human Umbilical Vein Endothelial Cells
IL-1β: Interleukin-1 beta
IL-4: Interleukin-4
IFN-γ: Interferon-γ
MSNs: Mesoporous silica nanoparticles
MFI: Mean fluorescence intensities
NPs: Nanoparticles
OVA: Ovalbumin
OD: Optical density
PAMPs: Pathogen-associated molecular patterns
PRRs: pattern recognition receptors
PBS: Phosphate buffered saline
PBST: PBS containing 0.1% Tween 20
RT: Room temperature
TLR: Toll-like receptor
TMB: Tetramethylbenzidine
VLPSi: Virus-like porous silica;

## Notes

### Competing Interest Statement

The authors have declared no competing interest.

## References

(1) Poria, R.; Kala, D.; Nagraik, R.; Dhir, Y.; Dhir, S.; Singh, B.; Kaushik, N. K.; Noorani, M. S.; Kaushal, A.; Gupta, S. Vaccine development: Current trends and technologies. Life Sci 2024, 336, 122331. DOI: 10.1016/j.lfs.2023.122331 From NLM Medline.

(2) Hafner, A. M.; Corthesy, B.; Merkle, H. P. Particulate formulations for the delivery of poly(I:C) as vaccine adjuvant. Adv Drug Deliv Rev 2013, 65 (10), 1386–1399. DOI: 10.1016/j.addr.2013.05.013 From NLM Medline.

(3) Zhao, T.; Cai, Y.; Jiang, Y.; He, X.; Wei, Y.; Yu, Y.; Tian, X. Vaccine adjuvants: mechanisms and platforms. Signal Transduct Target Ther 2023, 8 (1), 283. DOI: 10.1038/s41392-023-01557-7 From NLM Medline.

(4) Wculek, S. K.; Cueto, F. J.; Mujal, A. M.; Melero, I.; Krummel, M. F.; Sancho, D. Dendritic cells in cancer immunology and immunotherapy. Nat Rev Immunol 2020, 20 (1), 7–24. DOI: 10.1038/s41577-019-0210-z From NLM Medline.

(5) Ling, X.; Dong, Z.; He, J.; Chen, D.; He, D.; Guo, R.; He, Q.; Li, M. Advances in Polymer-Based Self-Adjuvanted Nanovaccines. Small 2025, 21 (16), e2409021. DOI: 10.1002/smll.202409021 From NLM Medline.

(6) Lyu, J. H.; Liou, G. G.; Wang, M.; Kan, M. C. The genetic, biophysical and immunological studies of a self-adjuvanted protein nanoparticle. Vaccine 2025, 56, 127087. DOI: 10.1016/j.vaccine.2025.127087 From NLM Medline.

(7) Liang, S.; Gao, S.; Fu, S.; Yuan, S.; Liu, J.; Liang, M.; Han, L.; Zhang, Z.; Liu, Y.; Zhang, N. Screening Natural Cholesterol Analogs to Assemble Self-Adjuvant Lipid Nanoparticles for Antigens Tagging Guided Therapeutic Tumor Vaccine. Adv Mater 2025, 37 (27), e2419182. DOI: 10.1002/adma.202419182 From NLM Medline.

(8) Dykman, L. A.; Staroverov, S. A.; Fomin, A. S.; Khanadeev, V. A.; Khlebtsov, B. N.; Bogatyrev, V. A. Gold nanoparticles as an adjuvant: Influence of size, shape, and technique of combination with CpG on antibody production. Int Immunopharmacol 2018, 54, 163–168. DOI: 10.1016/j.intimp.2017.11.008 From NLM Medline.

(9) Mahony, D.; Cavallaro, A. S.; Stahr, F.; Mahony, T. J.; Qiao, S. Z.; Mitter, N. Mesoporous silica nanoparticles act as a self-adjuvant for ovalbumin model antigen in mice. Small 2013, 9 (18), 3138–3146. DOI: 10.1002/smll.201300012 From NLM Medline.

(10) Balke, I.; Zeltins, A. Use of plant viruses and virus-like particles for the creation of novel vaccines. Adv Drug Deliv Rev 2019, 145, 119–129. DOI: 10.1016/j.addr.2018.08.007 From NLM Medline.

(11) Li, M.; Liang, Z.; Chen, C.; Yu, G.; Yao, Z.; Guo, Y.; Zhang, L.; Bao, H.; Fu, D.; Yang, X.;, et al. Virus-Like Particle-Templated Silica-Adjuvanted Nanovaccines with Enhanced Humoral and Cellular Immunity. ACS Nano 2022, 16 (7), 10482–10495. DOI: 10.1021/acsnano.2c01283 From NLM Medline.

(12) Hui, Y.; Yi, X.; Wibowo, D.; Yang, G.; Middelberg, A. P. J.; Gao, H.; Zhao, C. X. Nanoparticle elasticity regulates phagocytosis and cancer cell uptake. Sci Adv 2020, 6 (16), eaaz4316. DOI: 10.1126/sciadv.aaz4316 From NLM Medline.

(13) Christensen, D.; Henriksen-Lacey, M.; Kamath, A. T.; Lindenstrom, T.; Korsholm, K. S.; Christensen, J. P.; Rochat, A. F.; Lambert, P. H.; Andersen, P.; Siegrist, C. A.;, et al. A cationic vaccine adjuvant based on a saturated quaternary ammonium lipid have different in vivo distribution kinetics and display a distinct CD4 T cell-inducing capacity compared to its unsaturated analog. J Control Release 2012, 160 (3), 468–476. DOI: 10.1016/j.jconrel.2012.03.016 From NLM Medline.

(14) Wang, J.; Chen, H. J.; Hang, T.; Yu, Y.; Liu, G.; He, G.; Xiao, S.; Yang, B. R.; Yang, C.; Liu, F.;, et al. Physical activation of innate immunity by spiky particles. Nat Nanotechnol 2018, 13 (11), 1078–1086. DOI: 10.1038/s41565-018-0274-0 From NLM Medline.

(15) Wang, W.; Wang, P.; Tang, X.; Elzatahry, A. A.; Wang, S.; Al-Dahyan, D.; Zhao, M.; Yao, C.; Hung, C. T.; Zhu, X.;, et al. Facile Synthesis of Uniform Virus-like Mesoporous Silica Nanoparticles for Enhanced Cellular Internalization. ACS Cent Sci 2017, 3 (8), 839–846. DOI: 10.1021/acscentsci.7b00257 From NLM PubMed-not-MEDLINE.

(16) Wen, H.; Narvanen, A.; Jokivarsi, K.; Poutiainen, P.; Xu, W.; Lehto, V. P. A robust approach to make inorganic nanovectors biotraceable. Int J Pharm 2022, 624, 122040. DOI: 10.1016/j.ijpharm.2022.122040 From NLM Medline.

(17) Cooke, I. R.; Deserno, M. Solvent-free model for self-assembling fluid bilayer membranes: stabilization of the fluid phase based on broad attractive tail potentials. J Chem Phys 2005, 123 (22), 224710. DOI: 10.1063/1.2135785 From NLM Medline.

(18) Reynwar, B. J.; Illya, G.; Harmandaris, V. A.; Muller, M. M.; Kremer, K.; Deserno, M. Aggregation and vesiculation of membrane proteins by curvature-mediated interactions. Nature 2007, 447 (7143), 461–464. DOI: 10.1038/nature05840 From NLM Medline. Shi, X.; von dem Bussche, A.; Hurt, R. H.; Kane, A. B.; Gao, H. Cell entry of one-dimensional nanomaterials occurs by tip recognition and rotation. Nat Nanotechnol 2011, 6 (11), 714-719. DOI: 10.1038/nnano.2011.151 From NLM Medline. Sun, J.; Zhang, L.; Wang, J.; Feng, Q.; Liu, D.; Yin, Q.; Xu, D.; Wei, Y.; Ding, B.; Shi, X.; et al. Tunable rigidity of (polymeric core)-(lipid shell) nanoparticles for regulated cellular uptake. Adv Mater 2015, 27 (8), 1402-1407. DOI: 10.1002/adma.201404788 From NLM Medline.

(19) Li, J.; Dao, M.; Lim, C. T.; Suresh, S. Spectrin-level modeling of the cytoskeleton and optical tweezers stretching of the erythrocyte. Biophys J 2005, 88 (5), 3707–3719. DOI: 10.1529/biophysj.104.047332 From NLM Medline.

(20) Saric, A.; Cacciuto, A. Mechanism of membrane tube formation induced by adhesive nanocomponents. Phys Rev Lett 2012, 109 (18), 188101. DOI: 10.1103/PhysRevLett.109.188101 From NLM Medline. Wang, S.; Ma, S.; Li, H.; Dao, M.; Li, X.; Karniadakis, G. E. Two-component macrophage model for active phagocytosis with pseudopod formation. Biophys J 2024, 123 (9), 1069-1084. DOI: 10.1016/j.bpj.2024.03.026 From NLM Medline.

(21) Thompson, A. P.; Aktulga, H. M.; Berger, R.; Bolintineanu, D. S.; Brown, W. M.; Crozier, P. S.; in’t Veld, P. J.; Kohlmeyer, A.; Moore, S. G.; Nguyen, T. D.; et al. LAMMPS - a flexible simulation tool for particle-based materials modeling at the atomic, meso, and continuum scales. Computer Physics Communications 2022, 271, 108171. DOI: 10.1016/j.cpc.2021.108171.

(22) Berendsen, H.; Postma, J. P. M.; van Gunsteren, W.; DiNola, A. D.; Haak, J. R. Molecular-Dynamics with Coupling to An External Bath. The Journal of Chemical Physics 1984, 81, 3684. DOI: 10.1063/1.448118.

(23) Jarzynski, C. Nonequilibrium Equality for Free Energy Differences. Physical Review Letters 1997, 78 (14), 2690–2693. DOI: 10.1103/PhysRevLett.78.2690.

(24) Ma, S.; Qi, X.; Han, K.; Wang, S.; Hu, G.; Li, X. Computational investigation of flow dynamics and mechanical retention of age-associated red blood cells in the spleen. Physical Review Fluids 2023, 8 (6), 063103. DOI: 10.1103/PhysRevFluids.8.063103. Pivkin, I. V.; Peng, Z.; Karniadakis, G. E.; Buffet, P. A.; Dao, M.; Suresh, S. Biomechanics of red blood cells in human spleen and consequences for physiology and disease. *Proc Natl Acad Sci U S A* 2016, *113* (28), 7804-7809. DOI: 10.1073/pnas.1606751113 From NLM Medline.

(25) Tamarov, K.; Wang, J. T.; Kari, J.; Happonen, E.; Vesavaara, I.; Niemela, M.; Peramaki, P.; Al-Jamal, K. T.; Xu, W.; Lehto, V. P. Comparison between Fluorescence Imaging and Elemental Analysis to Determine Biodistribution of Inorganic Nanoparticles with Strong Light Absorption. ACS Appl Mater Interfaces 2021, 13 (34), 40392–40400. DOI: 10.1021/acsami.1c11875 From NLM Medline.

(26) Xu, W.; Pang, C.; Song, C.; Qian, J.; Feola, S.; Cerullo, V.; Fan, L.; Yu, H.; Lehto, V. P. Black porous silicon as a photothermal agent and immunoadjuvant for efficient antitumor immunotherapy. Acta Biomater 2022, 152, 473–483. DOI: 10.1016/j.actbio.2022.08.073 From NLM Medline.

(27) Hughes, C. S.; Foehr, S.; Garfield, D. A.; Furlong, E. E.; Steinmetz, L. M.; Krijgsveld, J. Ultrasensitive proteome analysis using paramagnetic bead technology. Mol Syst Biol 2014, 10 (10), 757. DOI: 10.15252/msb.20145625 From NLM Medline. Montaser, A. B.; Gao, F.; Peters, D.; Vainionpaa, K.; Zhibin, N.; Skowronska-Krawczyk, D.; Figeys, D.; Palczewski, K.; Leinonen, H. Retinal Proteome Profiling of Inherited Retinal Degeneration Across Three Different Mouse Models Suggests Common Drug Targets in Retinitis Pigmentosa. *Mol Cell Proteomics* 2024, *23* (11), 100855. DOI: 10.1016/j.mcpro.2024.100855 From NLM Medline.

(28) Demichev, V.; Messner, C. B.; Vernardis, S. I.; Lilley, K. S.; Ralser, M. DIA-NN: neural networks and interference correction enable deep proteome coverage in high throughput. Nat Methods 2020, 17 (1), 41–44. DOI: 10.1038/s41592-019-0638-x From NLM Medline.

(29) Zhou, Y.; Zhou, B.; Pache, L.; Chang, M.; Khodabakhshi, A. H.; Tanaseichuk, O.; Benner, C.; Chanda, S. K. Metascape provides a biologist-oriented resource for the analysis of systems-level datasets. Nat Commun 2019, 10 (1), 1523. DOI: 10.1038/s41467-019-09234-6 From NLM Medline.

(30) Briand, L.; Marcion, G.; Kriznik, A.; Heydel, J. M.; Artur, Y.; Garrido, C.; Seigneuric, R.; Neiers, F. A self-inducible heterologous protein expression system in Escherichia coli. Sci Rep 2016, 6, 33037. DOI: 10.1038/srep33037 From NLM Medline.

(31) Sun, G. G.; Wang, Z. Q.; Liu, C. Y.; Jiang, P.; Liu, R. D.; Wen, H.; Qi, X.; Wang, L.; Cui, J. Early serodiagnosis of trichinellosis by ELISA using excretory-secretory antigens of Trichinella spiralis adult worms. Parasit Vectors 2015, 8, 484. DOI: 10.1186/s13071-015-1094-9 From NLM Medline.

(32) Hartman, H.; Wang, Y.; Schroeder, H. W., Jr.; Cui, X. Absorbance summation: A novel approach for analyzing high-throughput ELISA data in the absence of a standard. PLoS One 2018, 13 (6), e0198528. DOI: 10.1371/journal.pone.0198528 From NLM Medline. Yu, X.; Gilbert, P. B.; Hioe, C. E.; Zolla-Pazner, S.; Self, S. G. Statistical approaches to analyzing HIV-1 neutralizing antibody assay data. *Stat Biopharm Res* 2012, *4* (1), 1-13. DOI: 10.1080/19466315.2011.633860 From NLM PubMed-not-MEDLINE.

(33) Chen, S.; Saeed, A.; Liu, Q.; Jiang, Q.; Xu, H.; Xiao, G. G.; Rao, L.; Duo, Y. Macrophages in immunoregulation and therapeutics. Signal Transduct Target Ther 2023, 8 (1), 207. DOI: 10.1038/s41392-023-01452-1 From NLM Medline. Racioppi, L.; Noeldner, P. K.; Lin, F.; Arvai, S.; Means, A. R. Calcium/calmodulin-dependent protein kinase kinase 2 regulates macrophage-mediated inflammatory responses. *J Biol Chem* 2012, *287* (14), 11579-11591. DOI: 10.1074/jbc.M111.336032 From NLM Medline.

(34) Hailemichael, Y.; Johnson, D. H.; Abdel-Wahab, N.; Foo, W. C.; Bentebibel, S. E.; Daher, M.; Haymaker, C.; Wani, K.; Saberian, C.; Ogata, D.;, et al. Interleukin-6 blockade abrogates immunotherapy toxicity and promotes tumor immunity. Cancer Cell 2022, 40 (5), 509–523 e506. DOI: 10.1016/j.ccell.2022.04.004 From NLM Medline.

(35) Jin, X. H.; Zheng, L. L.; Song, M. R.; Xu, W. S.; Kou, Y. N.; Zhou, Y.; Zhang, L. W.; Zhu, Y. N.; Wan, B.; Wei, Z. Y.;, et al. A nano silicon adjuvant enhances inactivated transmissible gastroenteritis vaccine through activation the Toll-like receptors and promotes humoral and cellular immune responses. Nanomedicine 2018, 14 (4), 1201–1212. DOI: 10.1016/j.nano.2018.02.010 From NLM Medline.

(36) Scheiblhofer, S.; Machado, Y.; Feinle, A.; Thalhamer, J.; Husing, N.; Weiss, R. Potential of nanoparticles for allergen-specific immunotherapy - use of silica nanoparticles as vaccination platform. Expert Opin Drug Deliv 2016, 13 (12), 1777–1788. DOI: 10.1080/17425247.2016.1203898 From NLM Medline.

(37) Liu, Q.; Zhou, Y.; Li, M.; Zhao, L.; Ren, J.; Li, D.; Tan, Z.; Wang, K.; Li, H.; Hussain, M.;, et al. Polyethylenimine Hybrid Thin-Shell Hollow Mesoporous Silica Nanoparticles as Vaccine Self-Adjuvants for Cancer Immunotherapy. ACS Appl Mater Interfaces 2019, 11 (51), 47798–47809. DOI: 10.1021/acsami.9b19446 From NLM Medline.

(38) Phan, N. M.; Nguyen, T. L.; Choi, Y.; Mo, X. W.; Trinh, T. A.; Yi, G. R.; Kim, J. High Cellular Internalization of Virus-Like Mesoporous Silica Nanoparticles Enhances Adaptive Antigen-Specific Immune Responses against Cancer. ACS Appl Mater Interfaces 2024, 16 (35), 45917–45928. DOI: 10.1021/acsami.4c07106 From NLM Medline.

