## Supplementary material for "Tunable Rigid Spikes on Virus-Like Porous Silica Enable Mechanistically Controlled Nanovaccine Platforms": Table S1-2 and Figure S1-11,

**Table S1**. Efficiency of NPs loading OVA protein

| Groups | Concentration of OVA (μg/mL) | Load efficiency |
|---|---|---|
| VLPSi-30-OVA | 164.94 | 8.25% |
| VLPSi-5-OVA | 156.94 | 7.85% |

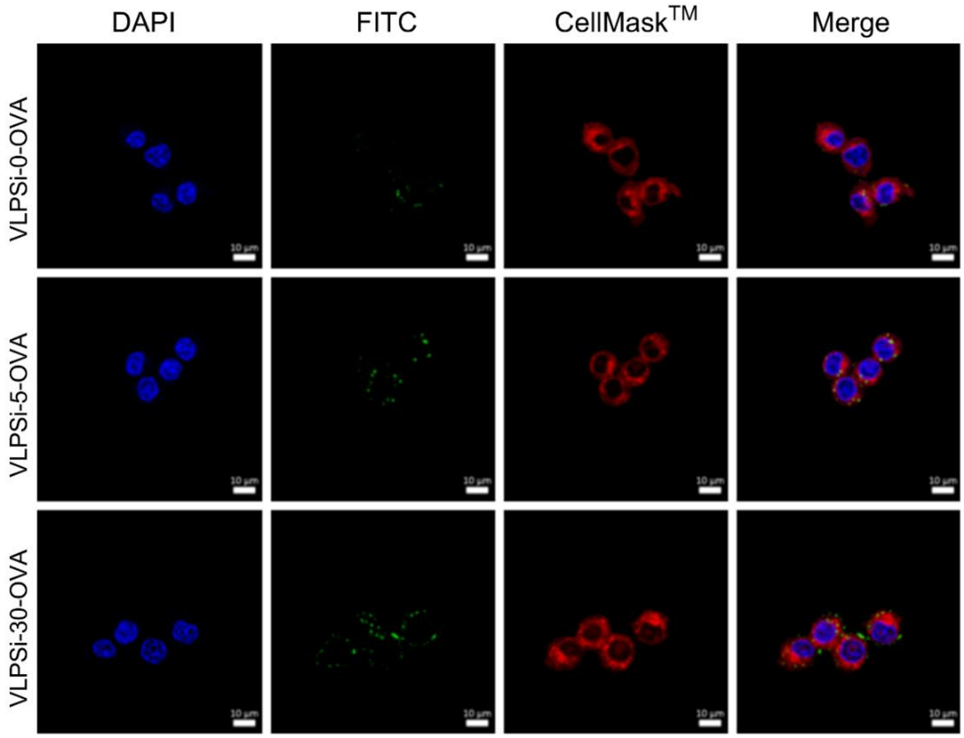


**Figure S1**. Detailed Confocal microscopy images of OVA-loaded VLPSi uptake with varying spike lengths by RAW264.7 cells. The VLPSi were labeled with FITC and incubated with the cells for 4 hours, with the fluorescence signal shown in green. The nucleus and cell membrane were stained with DAPI (blue) and CellMaskTM (red). The scale bar represents 10 μm.

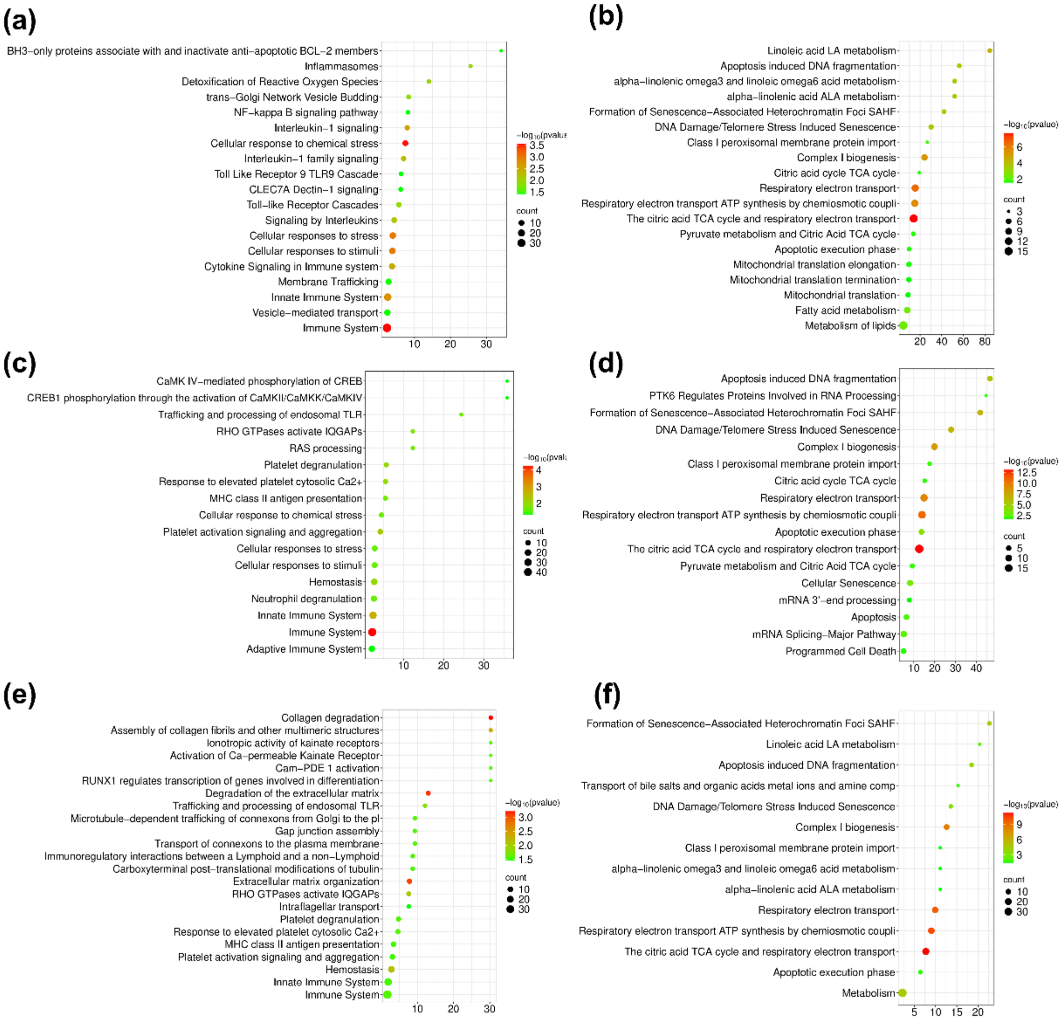


**Figure S2**. Pathway enrichment analysis of differentially expressed genes from Reactome database. (a, b) the pathway related to up-regulated and down-regulated genes of VLPSi-0-OVA group. (c, d) the pathway related to up-regulated and down-regulated genes of VLPSi-5-OVA group. (e, f) the pathway related to up-regulated and down-regulated genes of VLPSi-30-OVA group. Count indicates the number of genes (the bigger dots refer to larger amounts). The Fold Enrichment of a pathway is defined as the ratio of the percentage of differentially expressed genes to the corresponding percentage in the total gene amount. The horizontal axis represents gene enrichment, while the vertical axis represents the enriched pathway name. The dot color indicates different thresholds of the p-value.

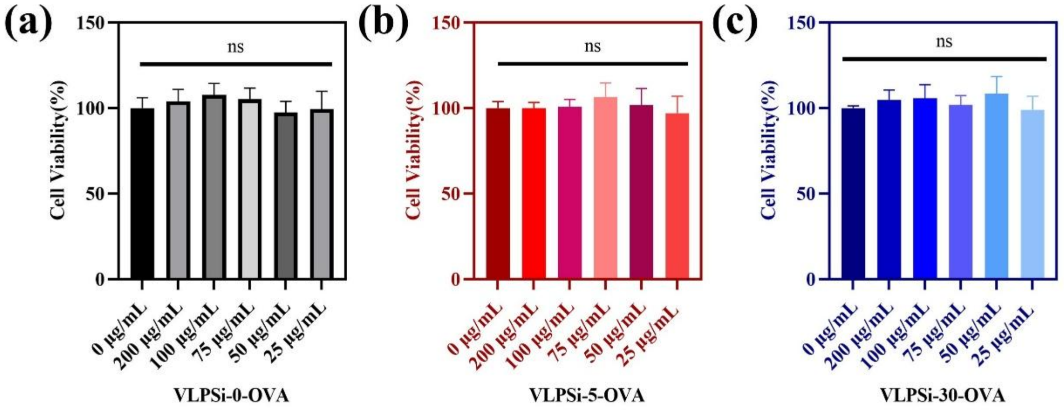


**Figure S3**. Figures a, b, c shows the toxicity of NPs-OVA on HUVEC cells;

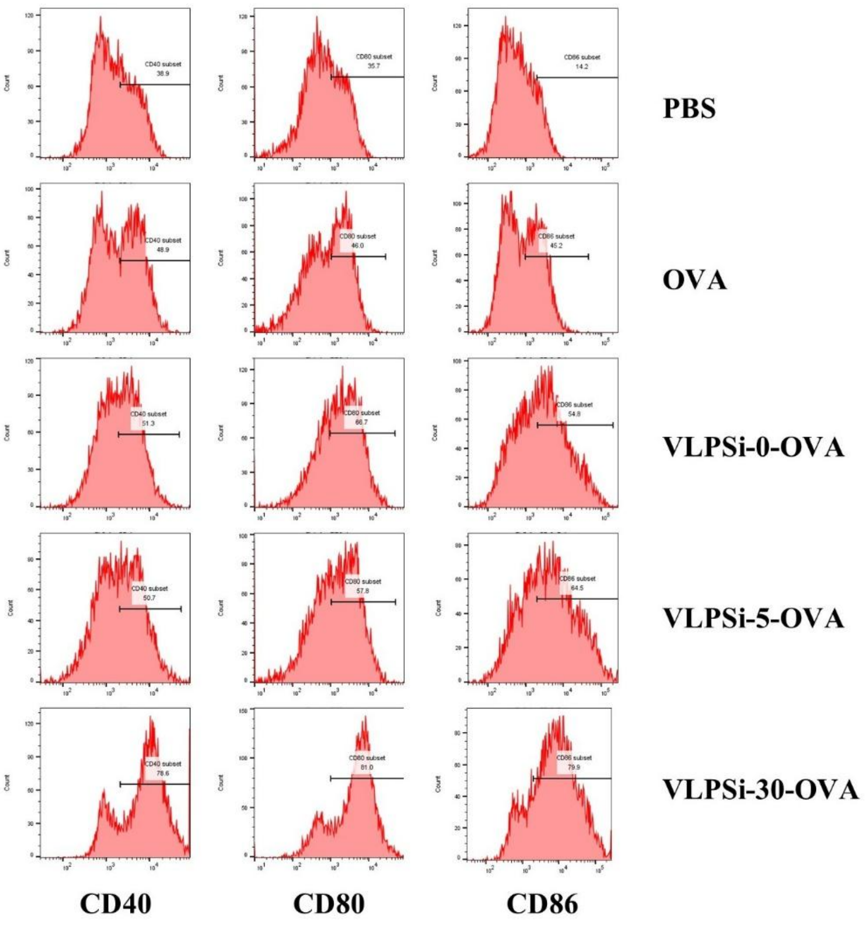


**Figure S4**. Effect of VLPSis-OVA on BMDC maturation

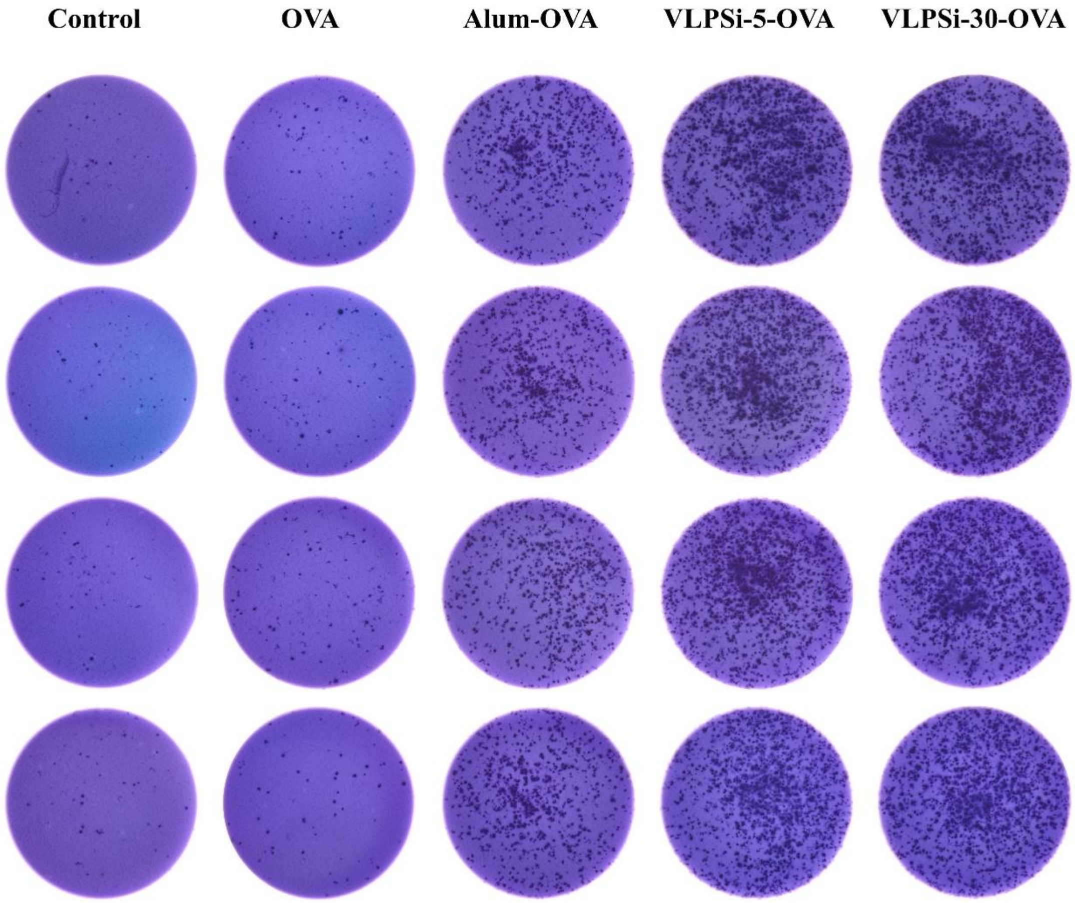


**Figure S5**. ELIspot results showing the effect of each group on the number of IL-4-secreting splenocytes in mice

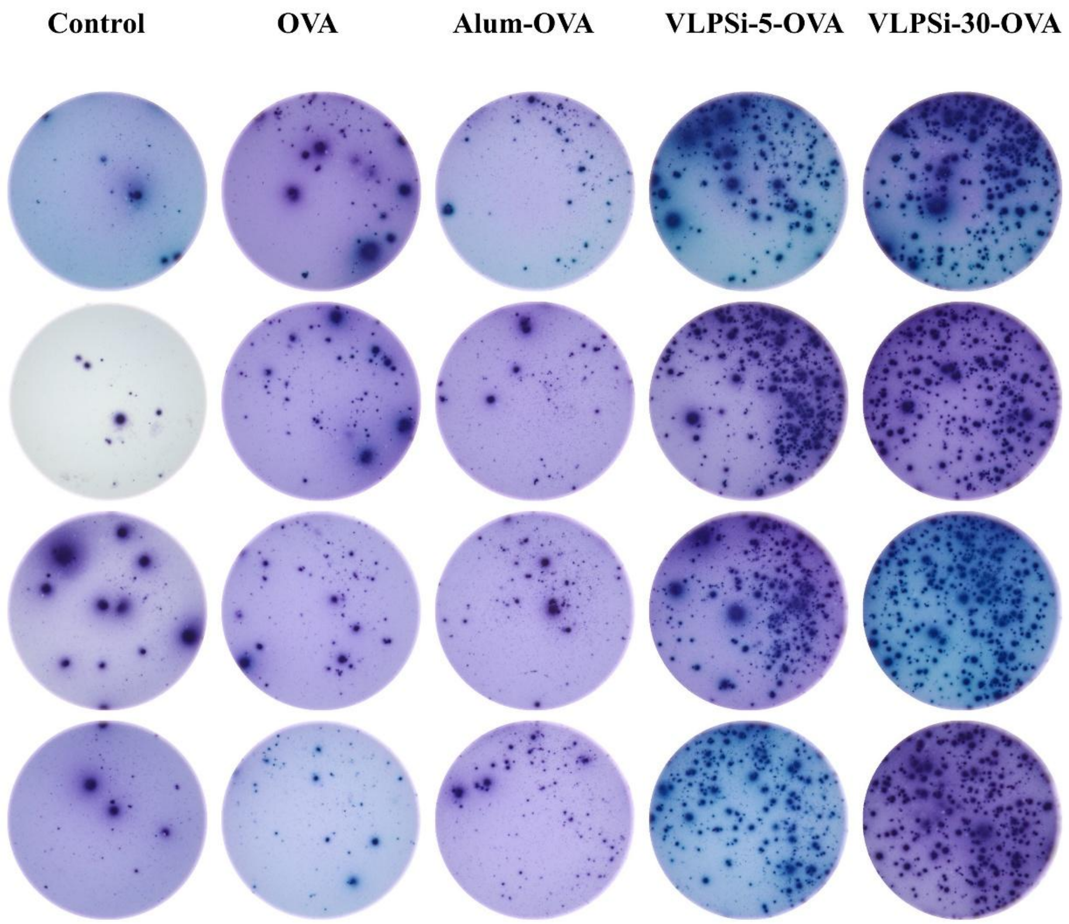


**Figure S6**. ELIspot results showing the effect of each group on the number of IFN-r-secreting splenocytes in mice

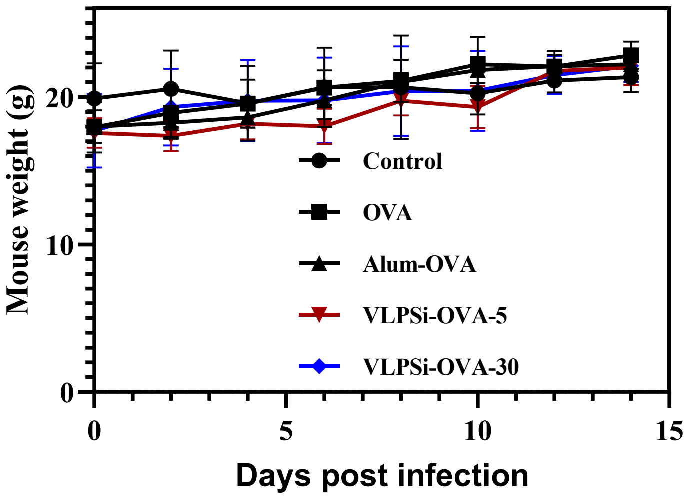


**Figure S7**. The changes in body weight of mice in each group.

**Table S2**. Efficiency of NPs loading rEsxB protein

| Groups | O.D. | Concentration of rEsxB (μg/mL) | Load efficiency |
|---|---|---|---|
| VLPSi-30-OVA | 0.4347 | 297.19 | 7.13% |
| VLPSi-5-OVA | 0.457 | 325.27 | 7.80% |

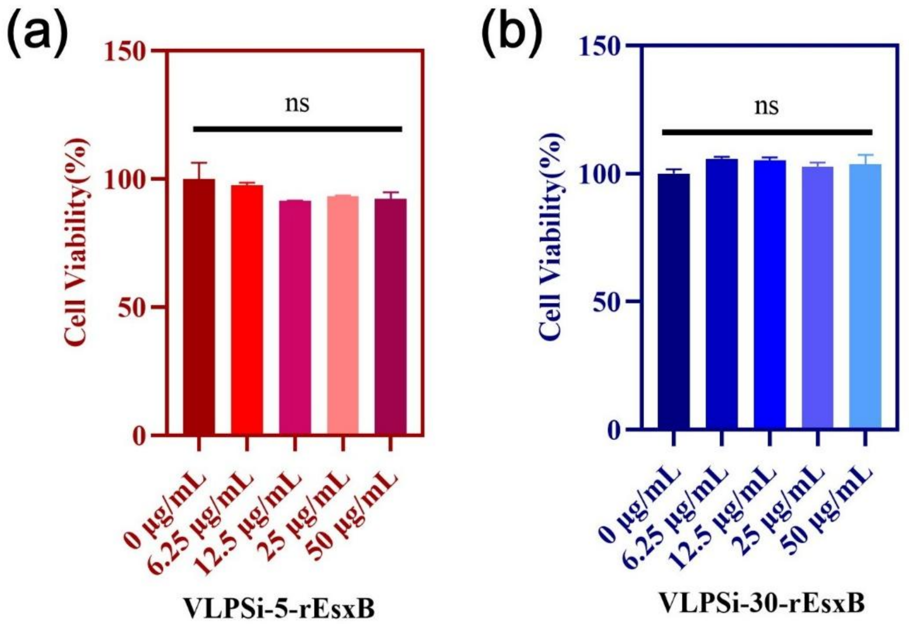


**Figure S8**. Figures a, b shows the cytotoxicity of NPs-rEsxB on BMDC cells.

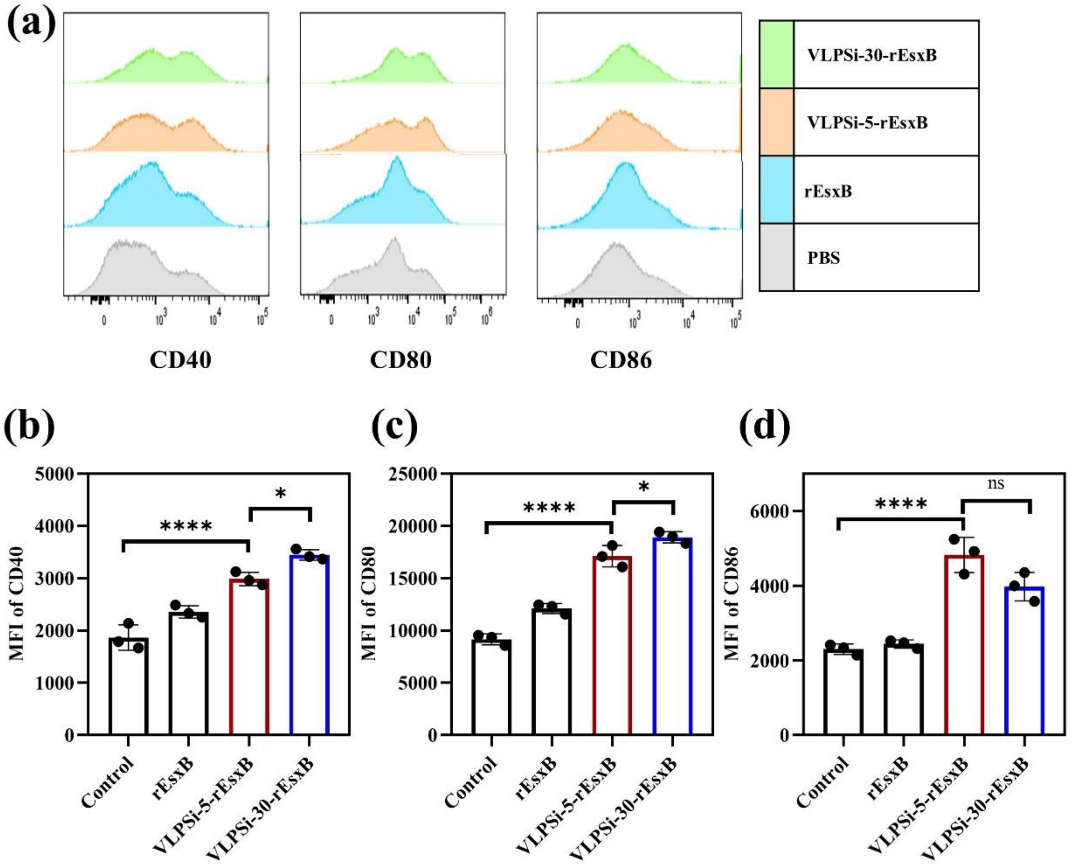


**Figure S9**. The effect of VLPSi-rEsxB on BMDC maturation. The P-value of less than 0.05 was considered statistically significant (* $p < 0.05$; ** $p < 0.01$; *** $p < 0.001$; **** $p < 0.0001$).

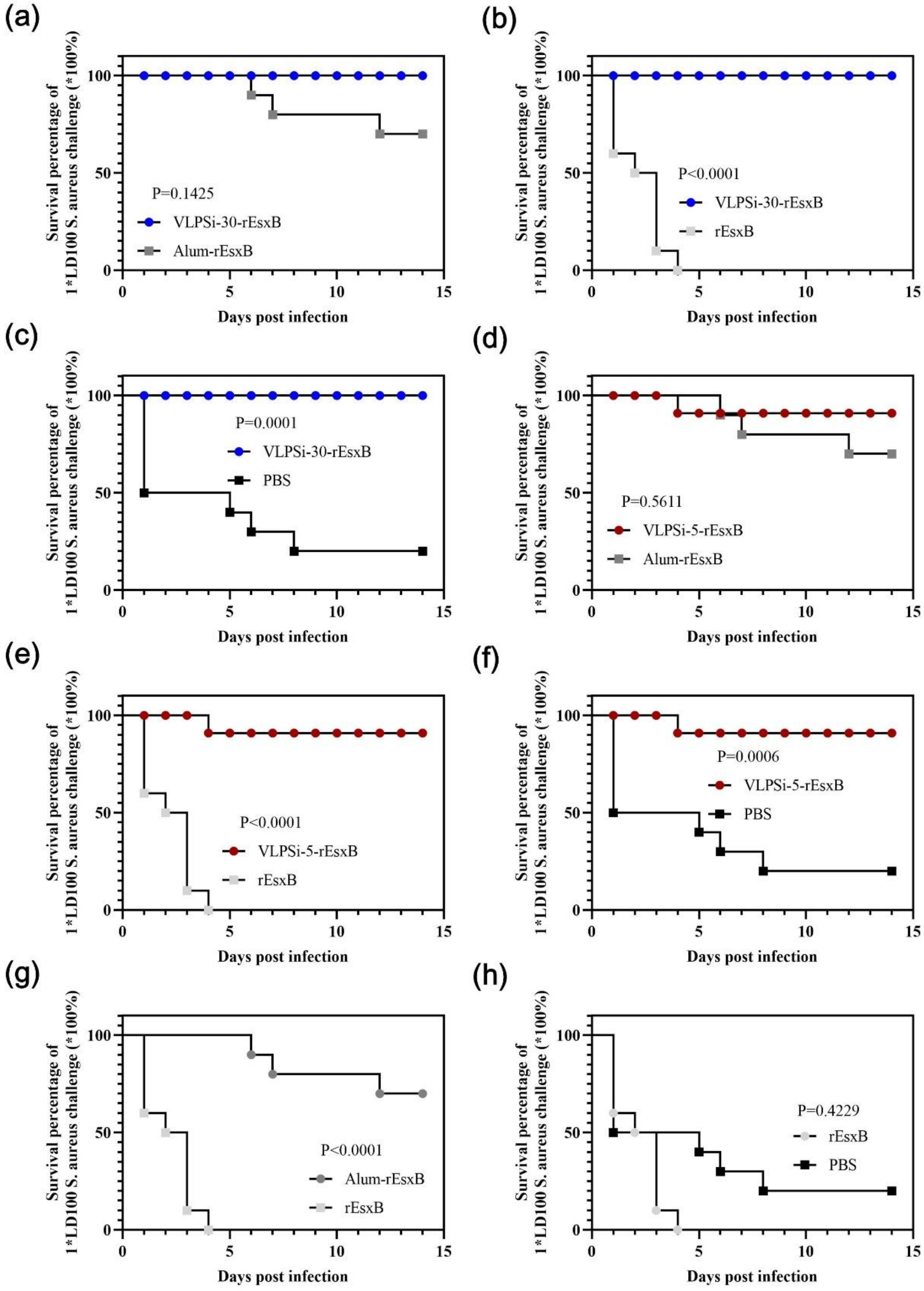


**Figure S10**. The survival rate of 1×LD100 S. aureus challenge experiment

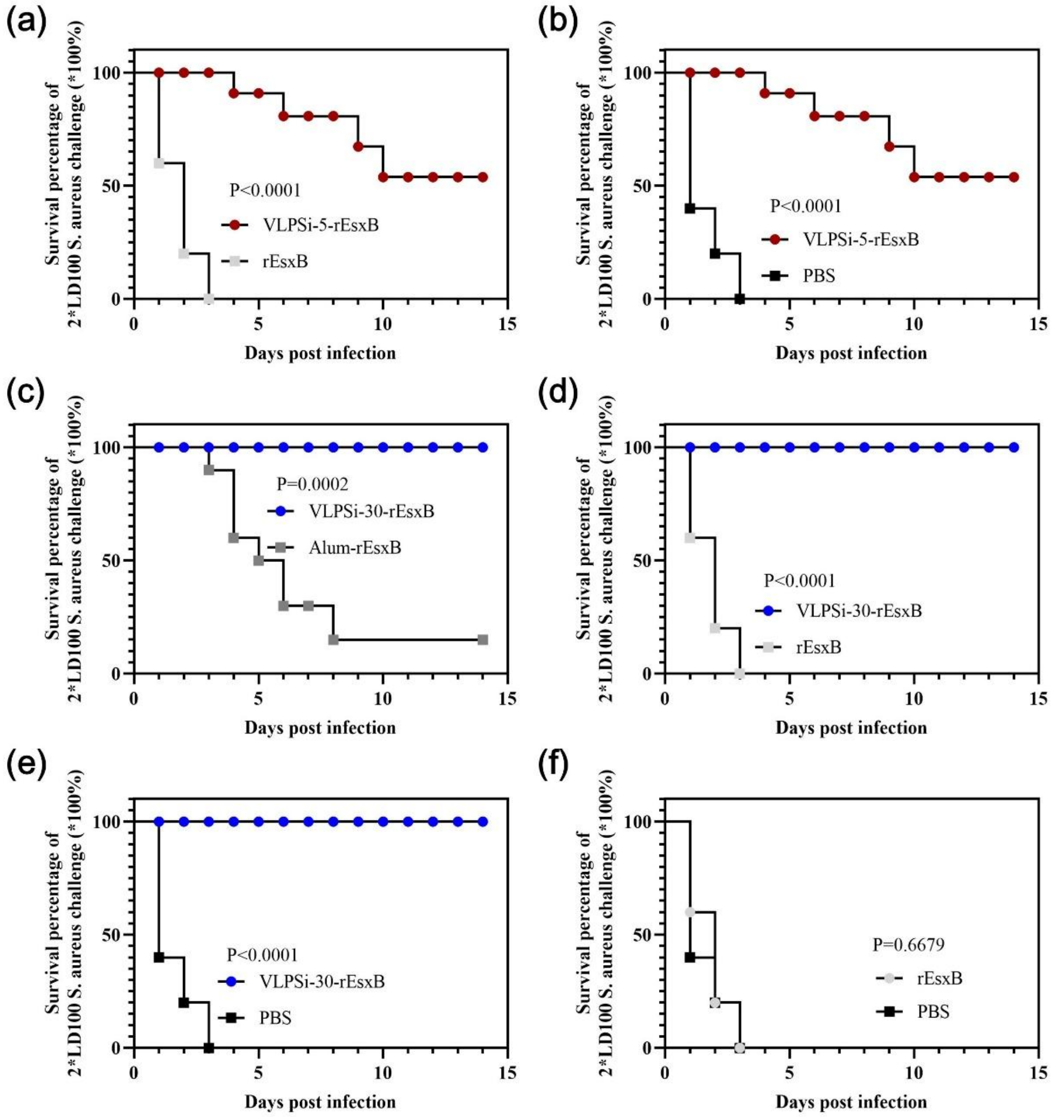


**Figure S11**. The survival rate of 2×LD100 S. aureus challenge experiment
